# Trade-off between light deprivation and desiccation in intertidal seagrasses due to periodic tidal inundation and exposure: insights from a data-calibrated model

**DOI:** 10.1101/2023.12.06.570383

**Authors:** Xinyan Wang, Matthew P. Adams, Dongdong Shao

## Abstract

Many seagrass species thrive in shallow intertidal zones globally, adapting to periodic tidal inundation and exposure with distinctive physiological traits and offering crucial ecosystem services. However, predicting the responses of intertidal seagrasses to external stressors is hampered by the complexity of the dynamic and harsh environments they occupy. Consequently, intertidal seagrass growth models, especially those incorporating dynamic physiological responses, are scarce in the literature. Our study comprehensively collated relevant data from the literature to parameterize the relationship between air exposure, seagrass leaf water content and photosynthetic efficiency to inform new growth rate functions for generalisable intertidal seagrass growth models. We tested the applicability of these model formulations for scenarios with varying physiological process assumptions, seagrass species, tidal conditions, meadow elevations and water turbidity. We found that neglecting air-exposed physiological responses (i.e., leaf water content loss and reduced photosynthetic efficiency) can substantially overestimate seagrass growth rates. We also observed a trade-off between light deprivation and desiccation on intertidal seagrass growth under specific tidal ranges and turbidity conditions. This can yield an “optimal” elevation where combined stressors of desiccation (increasing with meadow elevation) and light deprivation (decreasing with meadow elevation) are minimized. The predicted optimal elevation, i.e., the most suitable habitat for intertidal seagrass, moves upward as water turbidity increases. Our study provides both conceptual and quantitative guidance for ecological modellers to include air exposure responses of intertidal seagrasses in coastal ecosystem models. The model also helps to evaluate the viability of intertidal seagrass habitats and inform site selection for seagrass restoration.

## 1. Introduction

Seagrass meadows are among the most productive marine ecosystems in the world, and are widely distributed in both tropical and temperate coastal waters (Orth et al. 2020). Seagrasses are usually restricted by the upper depth limit due to air exposure at low tides and desiccation (Shafer et al. 2007; Suykerbuyk et al. 2018). As such, many seagrass species are intolerant to these conditions and are unable to grow in intertidal zones (Koch 2001). However, there are some species, including a few temperate species in the genus *Zostera* such as *Zostera japonica*, *Z. marina*, *Z. noltii*, as well as subtropical or tropical species such as *Halophila ovalis* and *Thalassia hemprichii*, that thrive in the shallow intertidal zones of estuaries, lagoons and other coastal areas (Colomer and Serra 2021; Shafer et al. 2007). Intertidal seagrass meadows function as essential foraging habitats (Espadero et al. 2020) and blue carbon stock, yielding higher organic carbon burial rates than subtidal seagrasses (de los Santos et al. 2022). In recent years, seagrass meadows have experienced continuous degradation caused by multiple stressors (Waycott et al. 2009). Meanwhile, intertidal seagrass ecosystems are subject to more dynamic and harsh environments, highlighting the complexity of assessing and predicting their dynamics and interactions with environmental stressors when aiming to inform their protection and restoration.

The clear dependence of seagrass growth on environmental conditions enables the development of mathematical models to represent the physiological relationships between environmental conditions and seagrasses (Scalpone et al. 2020). Mathematical models serve as useful tools for testing different environmental scenarios, offering insights that might not be achievable through traditional experiments. Existing seagrass models commonly include underwater light and temperature as driving forces on plant-scale processes including respiration, photosynthesis, and mortality (Elkalay et al. 2003; Piercy et al. 2023). Some models also include interspecific relationships such as the effects of algae and phytoplankton on light attenuation (Baird et al. 2016) and consumer-grazing effects (Turschwell et al. 2022). Other models expand their functionality by coupling with hydrodynamic and/or biogeochemical models to account for ecosystem-scale seagrass growth dynamics (Carr et al. 2012; Scalpone et al. 2020). However, the majority of the existing models lack the ability to simulate intertidal seagrass dynamics that are subject to periodic tidal inundation and air exposure (Erftemeijer et al. 2023; Folmer et al. 2012). To our best knowledge, only one previous study has developed formulations for the response of relative water content of intertidal seagrass leaves to different tidal conditions (Azevedo et al. 2017). However, it did not mathematically link the loss of relative water content to the photosynthetic process and subsequent vegetation growth dynamics. In the present work, we address this literature gap for the modelling of intertidal seagrasses.

Light deprivation due to tidal inundation is regarded as the most critical stressor influencing photosynthesis, growth and depth distribution of seagrasses including intertidal species (Bertelli and Unsworth 2018; Koch 2001). Alternatively, when exposed to air, intertidal seagrasses may exhibit photo-inhibition at high solar irradiances, and experience desiccation due to reduced leaf water content, both leading to declines in net photosynthesis (Kim et al. 2016). In addition, intertidal seagrasses may suffer from high temperatures when air-exposed, and the predominant impact of increased temperature during low tides is accelerated desiccation of seagrasses (Che et al. 2022). Conversely, many field studies have also found that intertidal seagrasses can avoid photo-inhibition due to their high tolerance to light stress without damage to photosynthetic apparatus (Clavier et al. 2011; Petrou et al. 2013). Consequently, intertidal seagrasses may take advantage of high irradiance during low tides which serve as a “window” of photosynthetic relief (Petrou et al. 2013). For example, a previous study has found that air-exposed *Z. noltii* in the south coast of Portugal exhibited increased productivity attributed to sustained leaf hydration (Silva et al. 2005). Yet, the same intertidal species *Z. noltii* in southern Spain showed a reduced photosynthesis rate attributed to severe desiccation during air exposure (Pérez-Lloréns et al. 1994). The contrasting results suggest that desiccation, which here refers to the reduction in the leaf water content, might be the key factor determining the photosynthetic responses of intertidal seagrasses. Therefore, light deprivation and desiccation are two dominant factors controlling intertidal seagrass growth. Light deprivation can only become potentially significant in the lower intertidal zone, whereas desiccation tends to have greater importance in the intermediate and upper zones (Cabaço et al. 2009). Since light deprivation effects on seagrasses have already been captured in the vast majority of seagrass models, we propose that physiological responses of intertidal seagrasses to air exposure, especially that triggered by alteration in leaf water content, should also be included in models of intertidal seagrass growth, to more accurately simulate their growth dynamics throughout tidal cycles.

New model formulations should ideally be informed by experimental data, and several experimental studies have examined the photosynthetic responses of intertidal seagrasses to air-exposed desiccation (Jiang et al. 2014; Leuschner et al. 1998; Shafer et al. 2007). Effective quantum yield of seagrass leaves is commonly measured as a metric of photosynthetic efficiency in these studies. Thus, the relative effect of air exposure on intertidal seagrass growth dynamics can be derived from the observed relationship between effective quantum yield and relative water content for different seagrass species.

Hence, the main objectives of this study are four-fold: 1) to comprehensively collate data from experimental studies to parameterize the relationship between air exposure, relative water content of seagrass leaves and photosynthetic efficiency; 2) to develop a generalisable intertidal seagrass growth model based on these relationships throughout the tidal cycles; 3) to provide illustrative parameterisations of this model for various physiological process assumptions, tidal conditions, meadow elevation, water column turbidity, and seagrass species; and 4) to examine how model predictions of seagrass growth rates are altered by the explicit inclusion of intertidal processes. Our study provides conceptual and quantitative guidance for ecological modellers who wish to include air exposure responses of intertidal seagrasses in their coastal ecosystem models. Ultimately, it is hoped that the improved modelling made possible from this work will assist in the evaluation of viable seagrass habitats for restoration activities.

## 2. Methods

### 2.1 Context for the intertidal seagrass model

We start by describing the mathematical modelling context to articulate the scientific gap that our intertidal seagrass model fills. As is typical in seagrass growth models, we assume that carbon accumulation is the rate-limiting step for plant growth (Moreno-Marín et al. 2018; Poorter et al. 2013). The net growth *dS/dt* of seagrasses is therefore assumed to be limited by the balance between carbon gain (photosynthesis, *P*) and carbon losses (e.g., respiration *R*, mortality *m* and dissolved organic carbon exudation *E*),

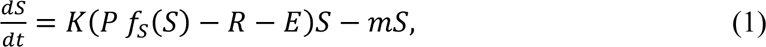

where *S* represents the local quantity of seagrass (typically written in units of dry-weight biomass per ground area, g DW m^-2^, or shoot density, shoots m^-2^), *t* is time (d), *K* is a conversion factor that accounts for the conversion of carbon gain/loss to seagrass gain/loss (e.g., in units of (g DW m^-2^)/(g C m^-2^) or (shoots m^-2^)/(g C m^-2^)), *P* is the gross photosynthesis rate (g C m^-2^ d^-1^), *f_s_*(S) is a crowding function (dimensionless) that limits the growth of seagrass at high densities due to self-shading, *R* is the respiration rate (g C m^-2^ d^-1^), *E* is the dissolved organic carbon exudation rate (g C m^-2^ d^-1^), and *m* is the mortality rate (d^-1^). It is common to define the crowding function so that *f_s_*(S) approximates unity at low density (i.e., at low values of *S*) and *f_s_*(S) decreases as *S* increases.

Eq. (1) is adapted from Kaldy (2012), and all loss processes described in Eq. (1) could be further split into losses from above-and below-ground biomass as needed. However, not all processes in Eq. (1) are included in all seagrass models; this equation is only provided here as a representative example of the modelling context in which our intertidal seagrass model is introduced (Section 2.2). In different models, seagrass *S* is quantified in either units of biomass (Jarvis et al. 2014; Turschwell et al. 2022) or shoot density (Adams et al. 2020; Carr et al. 2012), although these two quantities are positively correlated (Vieira et al. 2018). Similarly, different seagrass models assume different crowding functions *f_s_*(S), including the logistic growth function (Turschwell et al. 2022) or functions derived from geometric arguments (Baird et al. 2016); see Simpson et al. (2022) for other potentially relevant empirical crowding functions. For the remainder of this paper, it will neither be necessary to specify the units in which seagrass is quantified nor define the form of the crowding function; our model results will be equally applicable to all choices of seagrass density units and crowding function.

Gross photosynthesis is the only process in Eq. (1) contributing positively to seagrass growth. It is common to rewrite *KP* = *μ* so that Eq. (1) becomes

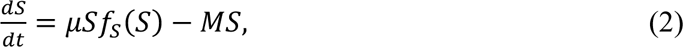

where *μ* is the per-capita growth rate (d^-1^) at low seagrass densities, and we have here grouped all loss terms into a per-capita loss rate *M* (d^-1^). In the present paper, we focus solely on the per-capita growth rate at low seagrass densities due to gross photosynthesis, *μ*.

Previous seagrass models assume that the growth rate *μ* (and by extension, the photosynthesis rate *P*) depends on light, temperature and/or nutrients (Baird et al. 2016; Elkalay et al. 2003; Turschwell et al. 2022). If information about the cumulative interaction between these factors is known (i.e., synergistic, additive or antagonistic), this information can be included in the mathematical definition of *μ* (Adams et al. 2020). However, in the absence of such information, two common methods of modelling the effects of interacting factors on growth rate are to assume a multiplicative (Turschwell et al. 2022) or the law of the minimum (Baird et al. 2016) formulation in the mathematical definition of *μ*. In the present work we will use both multiplicative (Eq. (3)) and law of the minimum (Eq. (4)) formulations of *μ*, and consider the effects of two controlling factors on *μ* that have particular relevance in the intertidal zone - light and relative water content (RWC) of seagrass leaves,

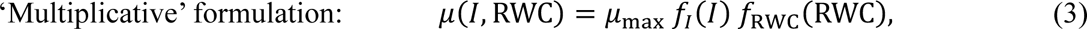

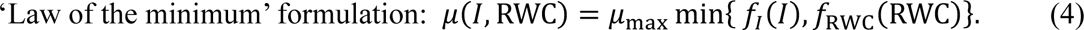

In these equations, *μ*_max_ is the maximum growth rate (d^-1^), *f_i_*(*I*) is a unitless function representing the effect of irradiance (i.e., light, or more precisely, photosynthetically active radiation or PAR) on the growth rate, and *f_RWC_*(RWC) is a unitless function representing the effect of RWC on the growth rate. Here, *I* (mol m^-2^ d^-1^) is the daily PAR dose at the seagrass canopy, and RWC is written as a fraction (bounded between 0 and 1 inclusive). It is assumed that the unitless functions *f_s_*(*I*) and *f_RWC_*(RWC) can only take values between 0 and 1 inclusive, so that the growth rate *μ* satisfies 0 ≤ *μ* ≤ *μ*_max_.

Although we are here only considering the effects of two controlling factors on the growth rate, each with its own unitless function in Eqs. (3) and (4), we point out that future applications of the model components we introduce could also incorporate other interacting factors (e.g., temperature, nutrients). These other factors could be incorporated in any subsequent models by the inclusion of appropriately defined additional functions within the multiplicative or law of the minimum formulation of the growth rate *μ* defined in Eqs. (3) and (4) respectively. We next describe each of the two unitless functions we focus on here (light and RWC) in further detail.

Various functional forms for the effect of irradiance on photosynthesis (and equivalently here, the growth rate) have been assumed in the literature (Jassby and Platt 1976); there is not yet standardisation of this functional form in the marine biological modelling community (Tian 2006). For the purposes of illustrating the new intertidal seagrass model components that we introduce in this paper, we chose the Michaelis-Menten function for

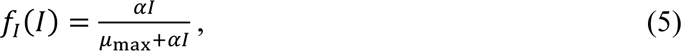

where *α* (d^-1^/(mol m^-2^ d^-1^)) indicates the efficiency of light utilisation for growth at low light.

### 2.2 Including intertidal effects: Modification of seagrass growth rate due to air-exposure

The photosynthesis rate of intertidal seagrasses is modified due to air exposure at low tides, and experimental data is available to parameterise this modification. Hence, we here describe mathematical relationships for how the seagrass growth rate may be altered by air exposure due to the loss of RWC in seagrass leaves when air exposed. The air-exposed responses of intertidal seagrasses are only triggered when the water depth drops to zero. The current conceptual understanding of the physiological processes for intertidal seagrasses when air-exposed and the associated equations considered in our model are shown in Fig. 1.

**Fig. 1.**
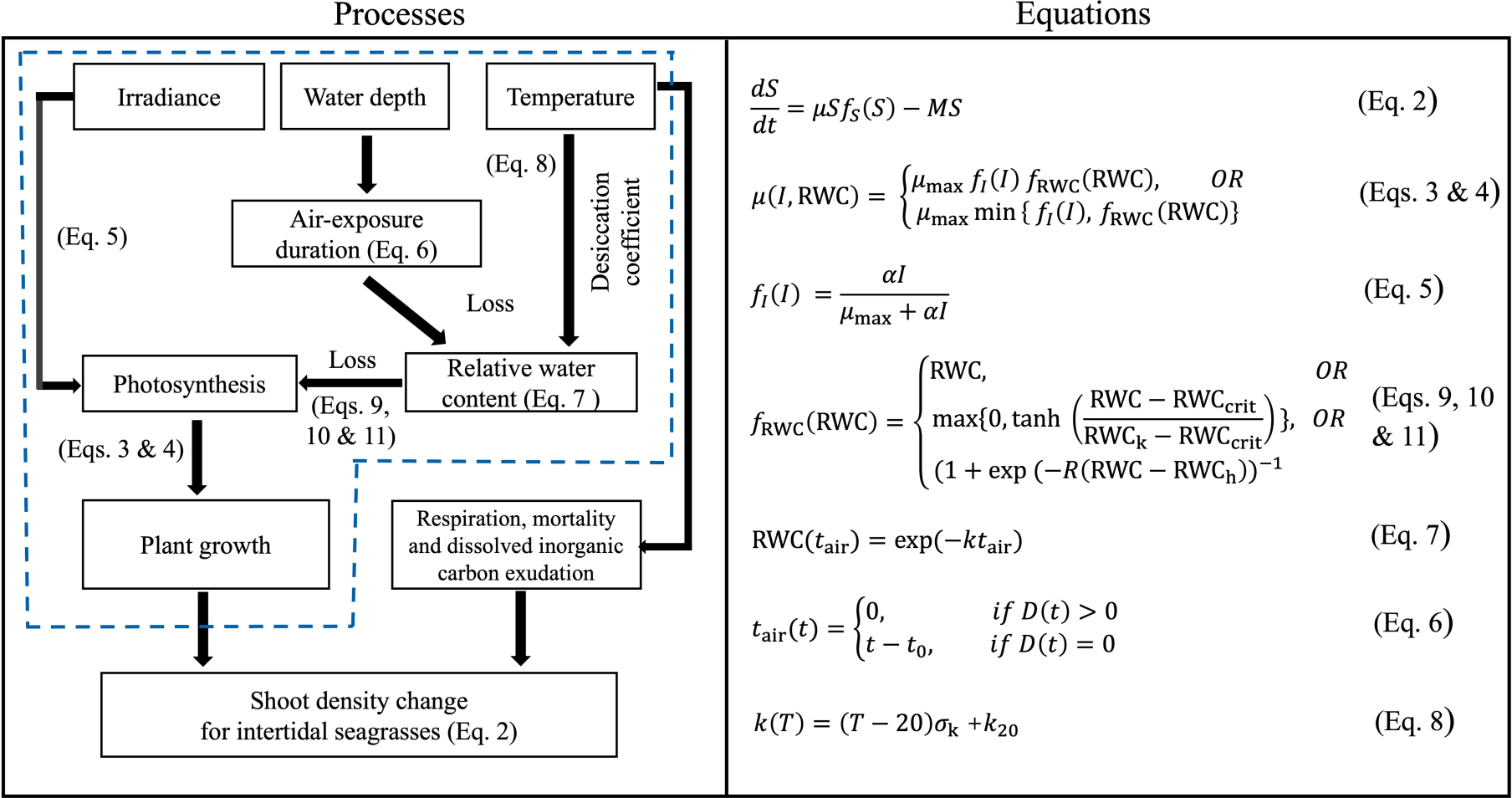
Schematic diagram of physiological processes and equations of intertidal seagrasses when air-exposed. The processes in the blue dashed box are quantitatively described in this study.

#### 2.2.1. The loss of relative water content

RWC of seagrass leaves decreases exponentially with time when seagrasses are exposed to air (Jiang et al. 2014; Papathanasiou et al. 2020; Shafer et al. 2007). For seagrasses present at the water depth *D* (m), the air-exposure duration *t_air_* can be defined as a function of time

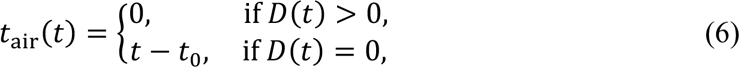

where *t*_0_ marks the time when the water depth *D* first drops to zero during a single exposure-inundation cycle. Here, when the water depth becomes zero (*D* = 0), it signifies the emersion (i.e., air exposure) of seagrasses, while a positive water depth (*D* > 0) indicates the inundation of the seagrasses. For clarity, Fig. 2 provides a visualization of how Eq. (6) represents the variation of air-exposure duration during a single exposure-inundation cycle.

**Fig. 2.**
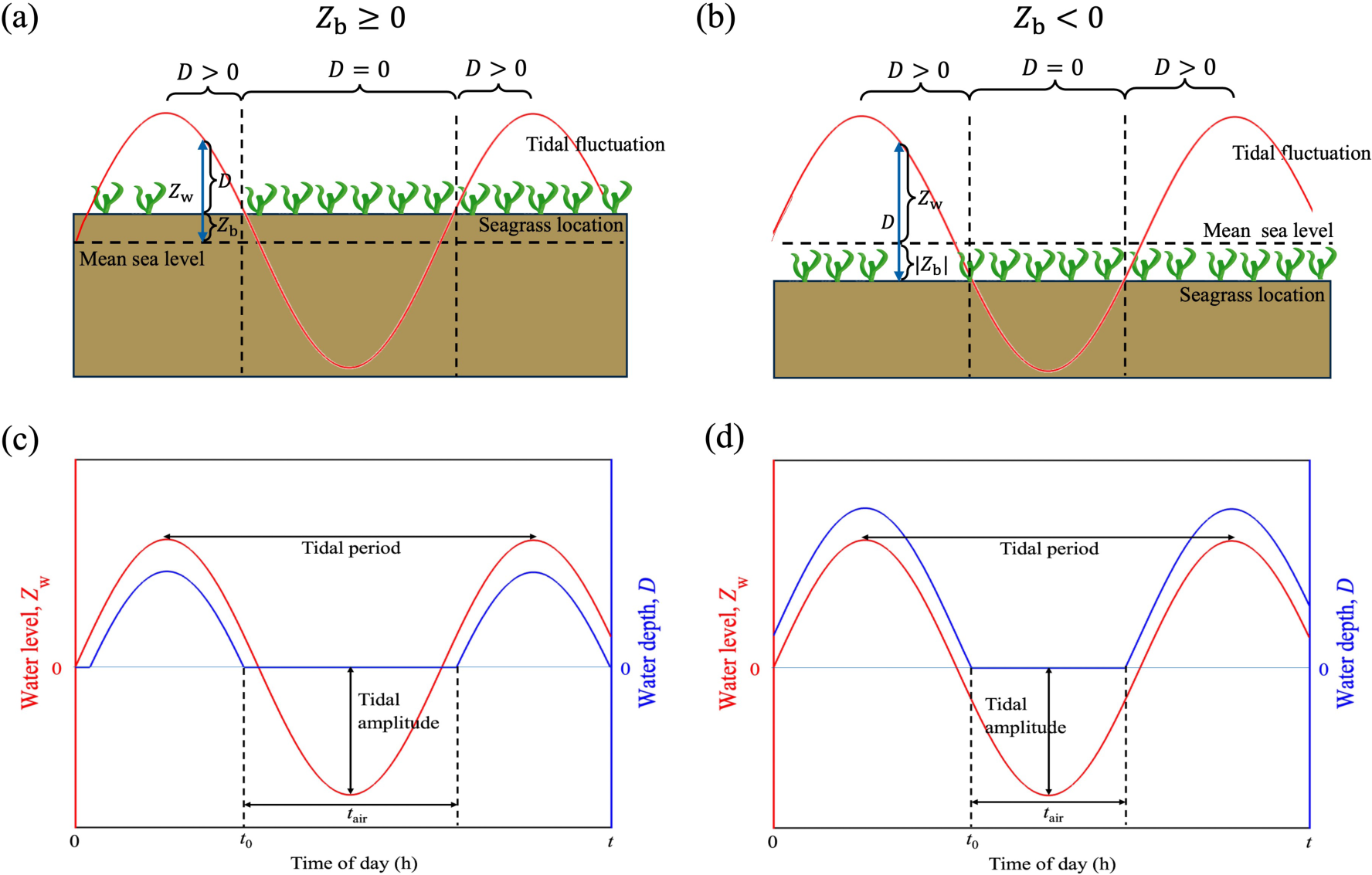
Variation of the water level *z*_w_ (*t*) and intertidal seagrass water depth *D*(*t*) throughout the tidal cycle. Seagrass meadows may grow at (a) an elevation that is above mean sea level, i.e., *z*_b_ ≥ 0, or (b) at an elevation that is below mean sea level, i.e., *z*_b_ < 0. In each of these two cases, the daily cycle of water level *z*_b_(*t*) and intertidal seagrass water depth *D*(*t*), can be assumed to approximately follow Eq. (12) and Eq. (13) respectively (introduced later in Section 2.3). The air-exposure duration *t*_&’(_ follows Eq. (6). For seagrass growing above mean sea level, this can result in daily time-series for *z*_b_(*t*) and *D*(*t*) following e.g., panel (c). For seagrass growing below mean sea level, this can result in daily time-series for *z*_b_(*t*) and *D*(*t*) following e.g., panel (d).

The exponential decline of RWC in seagrass leaves with air-exposure duration *t_air_* can be subsequently modelled as (Jiang et al. 2014; Seddon and Cheshire 2001; Shafer et al. 2007)

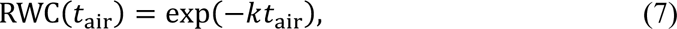

where *k* is the desiccation coefficient (units of d^-1^). As recovery of relative water content after re-submersion is expected to be relatively rapid (Azevedo et al. 2017), Eq. (7) inherently assumes that the fully hydrated state (i.e. RWC = 1) is instantly recovered for seagrass leaves after re-submersion. Thus, reduction in RWC only occurs when the water depth is zero (which corresponds to *t_air_* > 0, see Eq. (6)), otherwise the seagrass leaves remain fully hydrated (*t_air_* = 0 in Eq. (6)).

Experimentally, the desiccation coefficient (*k*) has been found to vary with different seagrass species and is positively correlated with air temperature *T* (Seddon and Cheshire 2001). Data for *k*(*T*) has thus far only been collected at a small number of air temperatures; in the absence of other data, this relationship is assumed to be linear,

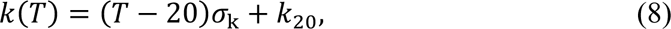

where σ_k_ (d^-1^ ℃^-1^) is the slope of the linear equation *k*(*T*) when fitted to data for desiccation coefficient *k* versus air temperature *T*, and *k*_20_ (d^-1^) is the desiccation coefficient at the air temperature of 20°C. If seagrass leaf desiccation data is not collected at different air temperatures, then Eq. (8) cannot be used. Conversely, this equation could be superseded by a more complicated (i.e., nonlinear) function if data for seagrass leaf desiccation coefficients is available for a large range of air temperatures.

As an example, we determined the parameters σ_k_ and *k*_20_ for two seagrass species (*Posidonia australis* and *Amphibolis antarctica*) from best-fit calibration of Eq. (7) and (8) to data from the laboratory desiccation experiment described in Seddon and Cheshire (2001). Their experiment measured how RWC varies with *t_air_* in *P. australis* and *A. antarctica* at four different air temperatures (18 °C, 24 °C, 28 °C, 32 °C). Figs. S1-S3 in the Supplementary Material show the plotted fits of Eq. (7) and (8) to the data, and Table S1 summarises the derived values of *k*(*T*), σ_k_ and *k*_20_ from these plotted fits. In Table 1, we compile data from Seddon and Cheshire (2001), as well as many other studies, to show the wide range of desiccation coefficients observed at different air temperatures across various seagrass species and locations. It is clear that the desiccation coefficients exhibit substantial variability, ranging from ∼ 8 d^-1^ to 260 d^34^.

**Table 1:**
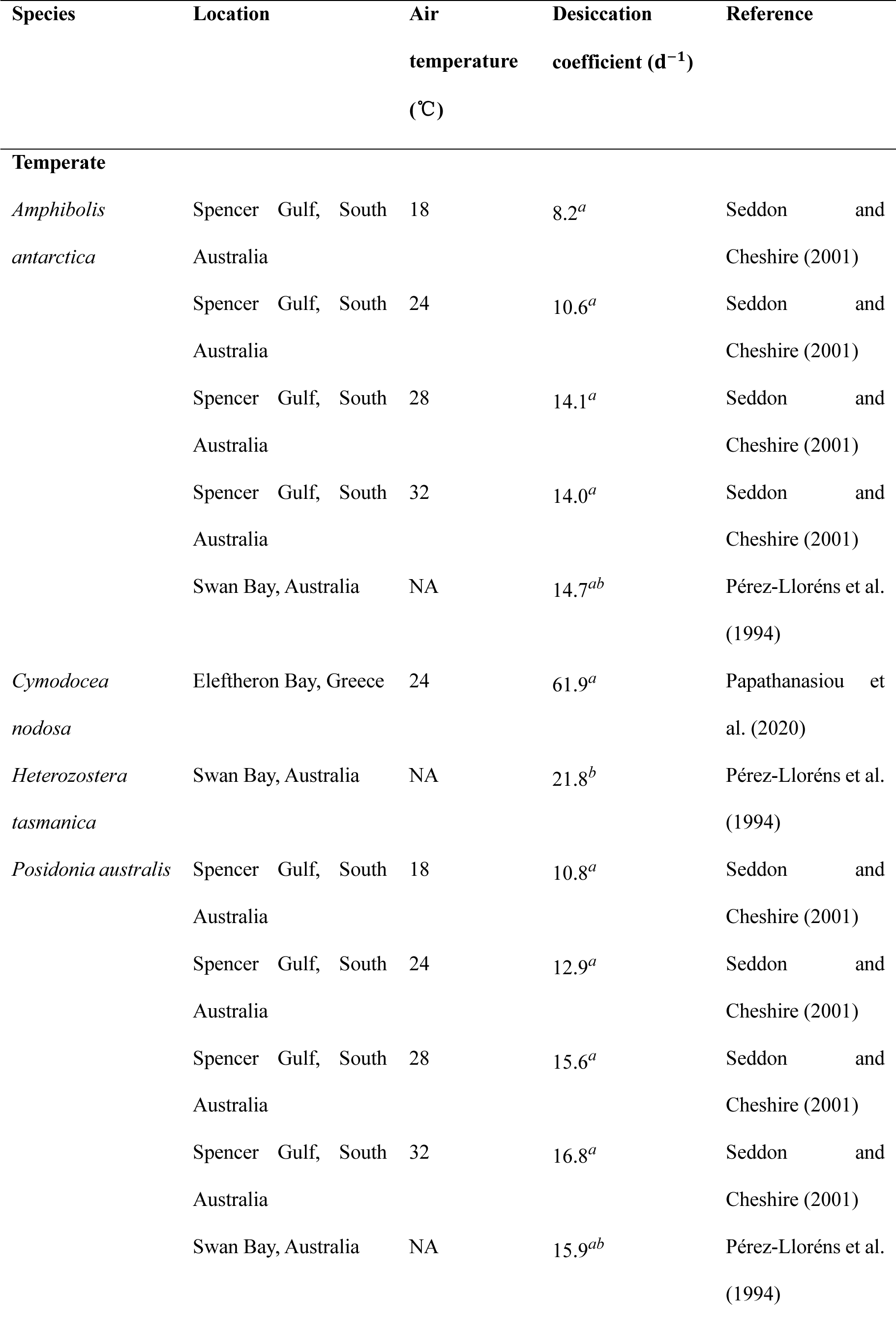
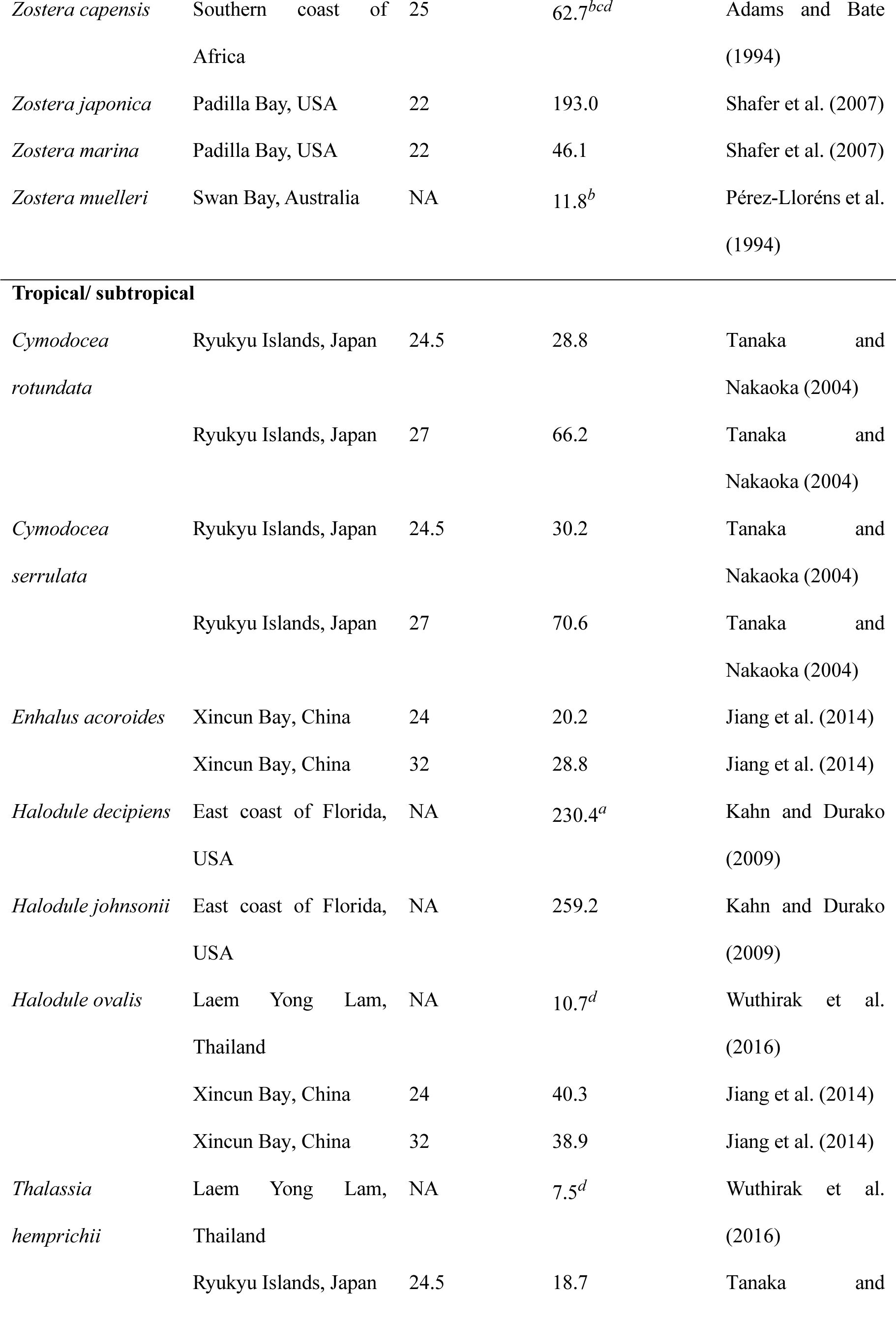
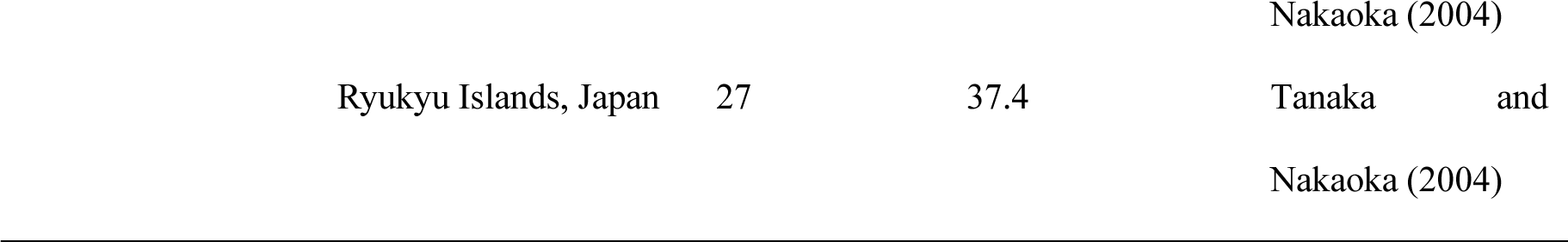
Summary of desiccation coefficients and corresponding air temperatures of intertidal/subtidal seagrasses from the literature.

#### 2.2.2. The response of photosynthetic efficiency to RWC loss

In Section 2.2.1 we described how RWC of seagrass leaves is altered during the tidal cycle, which is similar to the desiccation module described in Azevedo et al. (2017). Here, we go one step further and connect the RWC changes through to seagrass dynamics via the effect of RWC on seagrass photosynthesis rate.

Several experimental studies have reported a gradual decrease in effective quantum yield (a measure of photosynthetic efficiency) in seagrasses as RWC reduces (Jiang et al. 2014; Kahn and Durako 2009; Shafer et al. 2007). Here, we use these experimental findings to propose three alternative formulations for the unitless function *f_RWC_*(RWC) which describes the effect of RWC on seagrass growth rate (see Eqs. (3) and (4)). The precise form of the function *f_RWC_*(RWC) is presumed here to be species-specific due to the species-specific differences in their tolerance to desiccation, although they may be location-specific as well (Section 3.1). The three alternative forms of *f_RWC_*(RWC) that we propose are as follows:

**I. Linear model:** In Kahn and Durako (2009) and Papathanasiou et al. (2020), the relationship between effective quantum yield and relative water content (RWC) for two temperate intertidal species, *H. johnsonii* and *C. nodosa*, was linear (Supplementary Material Fig. S5). This suggests a linear form of *f_RWC_*(RWC) which we introduce as:

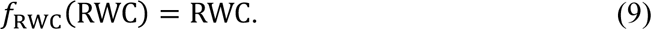

Eq. (9) gives that *f_RWC_*(RWC) = 1 when seagrass leaves are fully hydrated (RWC=1). The effective quantum yield is maximal, but usually less than one, when RWC=1 (Supplementary Material Fig. S5); this unavoidable inefficiency of the maximum effective quantum yield is scaled out in our mathematical formulation by the dimensionless form of *f_RWC_*(RWC).

**II. Hyperbolic tangent model:** In Jiang et al. (2014), the data for effective quantum yield versus RWC for tropical intertidal species *T. hemprichii* and *E. acoroides* were fitted to a standard hyperbolic tangent model. However, part of their data suggests that the value of *f_RWC_*(RWC) could equal zero for some range of RWC > 0 (see Supplementary Material Fig. S6), which is a behavior that is not possible using a standard hyperbolic tangent model unless a modification is made to its form. Thus, we introduced a modified hyperbolic tangent form of *f_RWC_*(RWC) to fit the data of Jiang et al. (2014); after scaling out the maximum effective quantum yield, this modified function *f_RWC_*(RWC) is:

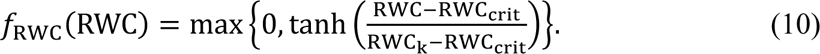

In Eq. (10), RWC_5/.6_ is the critical value of RWC below which the photosynthesis rate is zero, RWC_1_ is the value of RWC at which the photosynthesis rate is ∼76% of the maximum photosynthesis rate, and we enforce that the parameters RWC_5/.6_ and RWC_1_ must be non-negative. Mean parameter values for Eq. (10) fitted to *T. hemprichii* and *T. acoroides* data from Jiang et al. (2014) are provided in Supplementary Material Table S2.

**Ⅲ Sigmoidal curve model:** In Shafer et al. (2007), the data of effective quantum yield and RWC for temperate intertidal species *Z. japonica* and *Z. marina* were fitted to a sigmoidal curve model (see Supplementary Material Fig. S7). After scaling out the maximum effective quantum yield, this model for *f_RWC_*(RWC) can be written as:

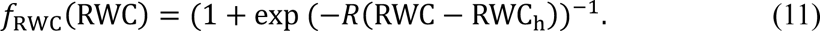

where *R* is the shape parameter of the fitted sigmoidal curve model, and RWC_7_ is the value of RWC that gives half of the maximum photosynthesis rate. Mean parameter values for Eq. (11) fitted to *Z. japonica* and *Z. marina* data from Shafer et al. (2007) are provided in Supplementary Material Table S2.

### 2.3 Simulating the intertidal cycle of air exposure and inundation

Now that a proposed modeling framework for intertidal seagrass has been fully described (Section 2.1, summarised in Fig. 1), we next sought to simulate models within this framework to explore what these models predict. This requires simulation of the intertidal seagrass response to dynamically changing environmental conditions, including daily fluctuations in water depth, irradiance and air temperature. Daily variation of environmental conditions can be quite complex; hence, it will be useful here to define “minimum realistic” models (Geary et al. 2020) of the external environmental conditions for the purposes of exploring the consequences of our intertidal seagrass model formulations. Here, we describe minimum realistic models for daily fluctuations in water depth, irradiance and air temperature conditions, and later in Section 2.4.2 we describe how we will use these models as environmental forcings for simulating our intertidal seagrass model.

First, the tidal cycles affect the water depth in the intertidal seagrass meadows, leading to periodic air exposure of seagrasses. We assumed that air exposure of seagrasses was forced by the M2 and S2 tidal constituents (i.e., the dominant constituents that lead to typical spring-neap tidal cycles) in the intertidal zones. Therefore, two superimposed cosine curves with different amplitudes and periods representing the M2 and S2 tidal constituents (Balke et al. 2016) were simulated to determine the water level relative to mean sea level,

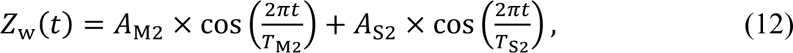

where *z*_w_(*t*) is the water level (m) relative to mean sea level at time *t* (d), *A*_M2_ is the amplitude of the M2 tidal constituent (m), *T*_M2_ is the period of the M2 tidal constituent (d), *A*_s2_ is the amplitude of the S2 tidal constituent (m), and *T*_<2_ is the period of the S2 tidal constituent (d). Consequently, the water depth *D* (m) at an intertidal seagrass meadow of interest is dictated by the changes in water level relative to meadow elevation,

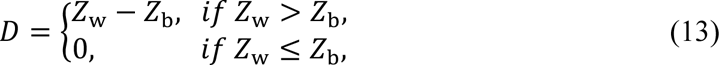

where *z*_=_ is the elevation of seagrass meadow relative to mean sea level (m). The relationship between water level *z*_w_ and intertidal seagrass water depth *D* throughout the tidal cycle is visualised in Fig. 2c & 2d.

Second, a minimum realistic model for daily light fluctuations is as follows. The within-daily water surface light *I*_>_(*t*) (in units of mol m^-2^ d^-1^) varies sinusoidally during the day, peaking at solar noon (i.e., 12 hours after solar midnight). However, the water surface light *I*_>_(*t*) must also be zero at night. This can be accomplished by using a sinusoidal curve for *I*_>_(*t*) that is truncated to be non-negative according to Johnson and Thornley (1984) and Adams et al. (2020),

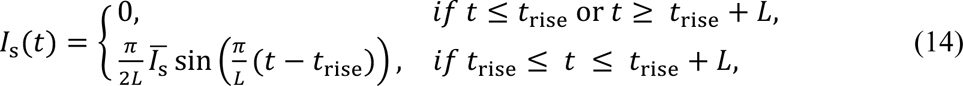

Where I*_S_* is the daily average water surface light (mol m^-2^ d^-1^), *t* is the time since solar midnight (d), *t*_/.>?_ is the daily sunrise time (d), and *L* is the day length (from sunrise to sunset) (d). Because tidal inundation of the seagrasses reduces the light they receive, the benthic light *I*(*t*) experienced by the seagrasses can then be calculated from the Beer-Lambert law (Kirk 1985),

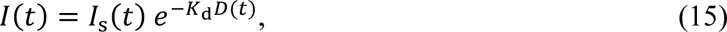

where *K*_d_ is the light attenuation coefficient in the water column (m^-1^), assumed here to be spatiotemporally constant for simplicity.

Finally, daily fluctuations in air temperature *T*_%./_(*t*) (in units of °C) can be coarsely approximated by a sinusoidal variation that peaks at a maximum air temperature some time *t*_∅_ (in d) after solar noon (Adams et al. 2020),

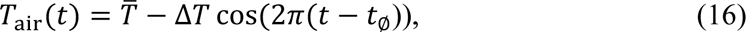

where *T* is the mean daily air temperature (℃) and Δ*T* (℃) is the maximum variation of daily air temperature from its mean value. A visualisation of Eqs. (14) and (16) is provided in Supplementary Material Fig. S8.

### 2.4 Model simulations

Using the data-calibrated intertidal seagrass model formulations and minimum realistic models for daily fluctuating environmental parameters introduced in the previous sections, we produced model simulations to explore both the applicability and consequences of the introduced formulations. First, we compared our data-calibrated formulations of seagrass photosynthetic efficiency reduction due to desiccation (captured by the function *f_s_*_RWC_(RWC)) to previously published experimental results for the same seagrass species at different locations and for the same and different tissue types (i.e., leaf versus shoot). Further explanation of this comparison is provided in Section 2.4.1. Second, seagrass growth rates were simulated under a variety of physiological process assumptions and environmental scenarios to identify generalisable conclusions obtained from our introduced intertidal seagrass model formulations (Section 2.4.2).

#### 2.4.1. Investigating the applicability of a parameterised intertidal seagrass model to other locations and other seagrass tissue types

We compared our data-calibrated formulations of seagrass photosynthetic efficiency reduction due to desiccation (*f_s_*_RWC_(RWC)) to data from additional experimental studies (Kim et al. 2020; Park et al. 2016). The purpose of this analysis was to identify the applicability of *f_s_*_RWC_(RWC) functions parameterised by data from a seagrass species at a particular location and with a particular tissue type, to the same species (1) in different locations, and (2) for the same and different tissue types. This is an important modelling question to consider, as environmental models are often parameterised using data from one location and applied to another location. We sought to understand the validity of such an application for our introduced intertidal seagrass model formulations.

To accomplish this, we simulated the relationship between photosynthetic efficiency scaled by its maximum, i.e., *f_RWC_*(RWC), and air-exposure duration *t_air_*, using Eqs. (7) and (11) parameterised to data (Supplementary Material Fig. S7) obtained for *Z. marina* and *Z. japonica* leaves in Padilla Bay, USA (Shafer et al. 2007). We compared these predictions of *f_RWC_*(RWC) to measured photosynthetic efficiency changes for *Z. marina* and *Z. japonica* reported in two experimental studies that were both carried out in the southern coast of South Korea (Kim et al. 2020; Park et al. 2016). Two datasets for *Z. marina* (Park et al. 2016) were available-both were for seagrass shoots, but at different sites within the southern coast of South Korea (Aenggang Bay and Koje Bay). Two datasets for *Z. japonica* (Kim et al. 2020) were also available – both were measured at Koje Bay, but for different seagrass tissues (leaves and shoots). Hence these datasets allowed us to examine the applicability of *f_RWC_*(RWC) formulations parameterised for seagrass species in one location to the same seagrass species in other locations and for the same and different seagrass tissues.

#### 2.4.2. Investigating the dependence of intertidal seagrass growth rate on model assumptions and environmental conditions

We then conducted a plethora of simulation scenarios for the introduced growth rate function *μ*(*I*, RWC) using Eqs. (6)-(11), with environmental forcings provided by daily fluctuations in water depth, light, and air temperature (Eqs. (12)-(16)). These scenarios allowed us to investigate (1) what effects do inclusion of seagrass physiological responses to intertidal processes have on their growth rates (i.e. including the factor *f_RWC_*(RWC) in seagrass models), (2) what differences in growth rates arise from different model assumptions (i.e., the assumption of which form of *f_s_*_()*_(RWC) (Eq. (9)-(11)); and the assumption of whether cumulative stressors interact multiplicatively (Eq. (3)) or if seagrass respond only to the strongest stressor (Eq. (4)), and (3) the effects of water turbidity, meadow elevation and tidal range on intertidal seagrass growth rate. For the remainder of this section, we detail what simulations were performed to undertake these investigations. In all simulations, the total period simulated was 15 d to cover an entire spring-neap cycle, and we assumed that the within-daily fluctuations of light and air temperature did not change from day to day.

**Baseline scenario.** We first describe a “Baseline” scenario for our simulations, which represents a specific set of environmental and seagrass growth characteristics that all our other “testing” scenarios are compared to. In the Baseline scenario, the cumulative effect of light deprivation and desiccation on seagrass growth *μ*(*I*, RWC) was assumed to follow the multiplicative formulation (Eq. (3)). In some of our testing scenarios described later, we also examined the law of the minimum formulation given in Eq. (4).

For the Baseline scenario, we parameterised the model simulations for the common intertidal seagrass genera *Zostera*, using parameters drawn (where possible) from data for intertidal *Z. japonica* meadows in the Yellow River Estuary (YRE), China. This choice of species and location for model parameterisation is relatively arbitrary since we are not particularly interested in the precise quantitative predictions of any individual simulation; instead, we are here primarily interested in comparing simulations between scenarios. *Z. japonica* is a reasonable species choice for this purpose because it is one of the most widely distributed seagrass species in China’s coastal areas. Similarly, the YRE is a reasonable location choice because it contains the largest habitat of *Z. japonica* in China (Zhou et al. 2022), where we have performed extensive field monitoring and experiments (Wang et al. 2022; Wang et al. 2021).

Hence, we drew parameters for the daily fluctuations of air temperature and light from a mixture of published (Zhang et al. 2019) and unpublished data for YRE (Table 2). Specifically, light and air temperature parameters were obtained from monitoring data from June to August in a typical growing season of the intertidal seagrass meadows in the YRE. Due to the lack of physiological data for *Z. japonica* available at YRE, seagrass photosynthesis responses to RWC were instead obtained from data for *Z. japonica* growing in the similar temperate region of Padilla Bay, USA (Shafer et al. 2007), for which the sigmoidal curve model for *f_RWC_*(RWC) given in Eq. (11) has already been fitted in the present study (Supplementary Material Table S2 and Supplementary Material Fig. S7). Similarly, we could not find a parameterisation of the photosynthesis-irradiance relationship for *Z. japonica*, and thus seagrass growth responses to light were parameterised from the related species *Z. marina* growing in Danish waters (Olesen and Sand-Jensen 1993) which possess a similar temperate climate to the YRE.

**Table 2:**
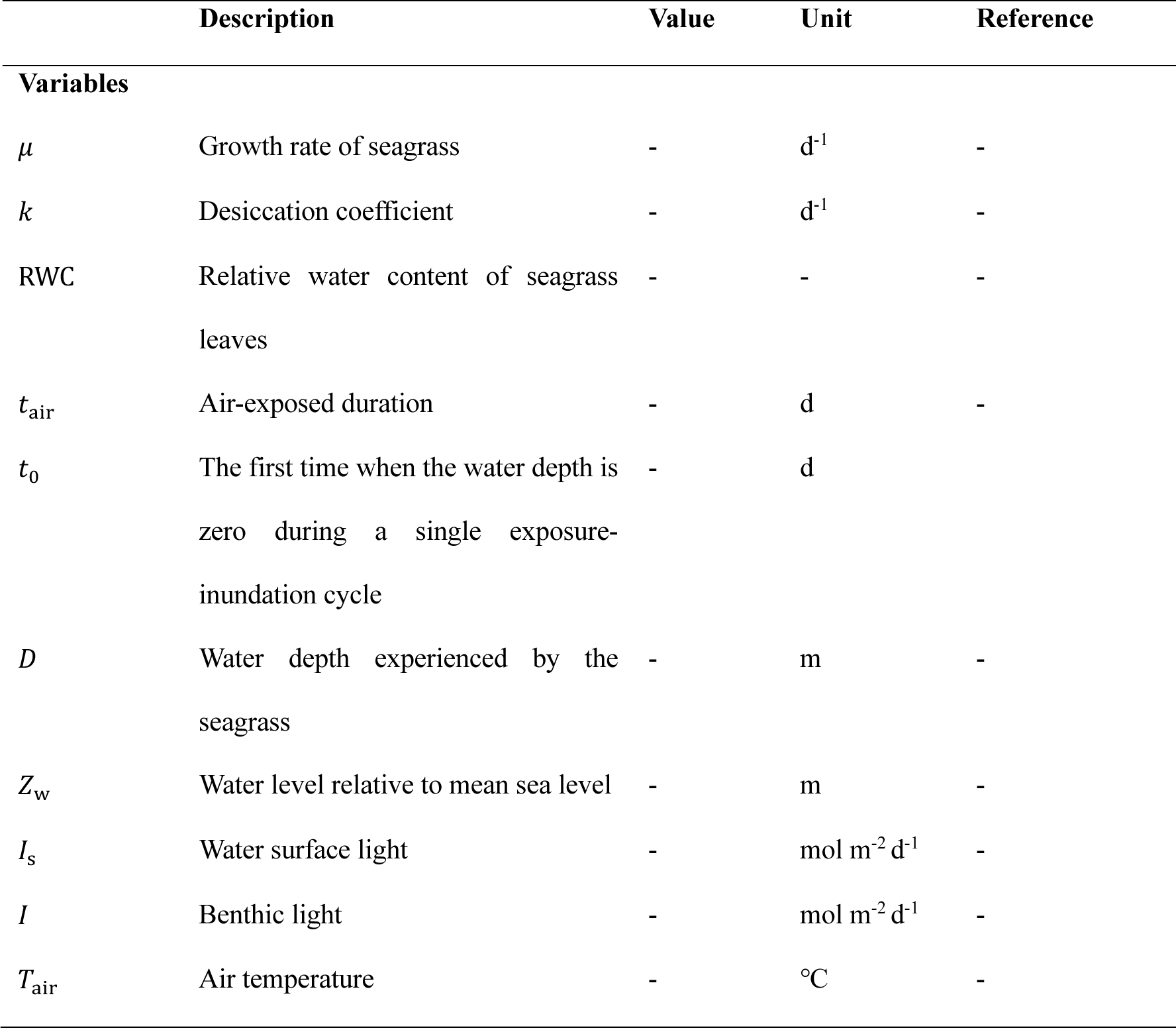
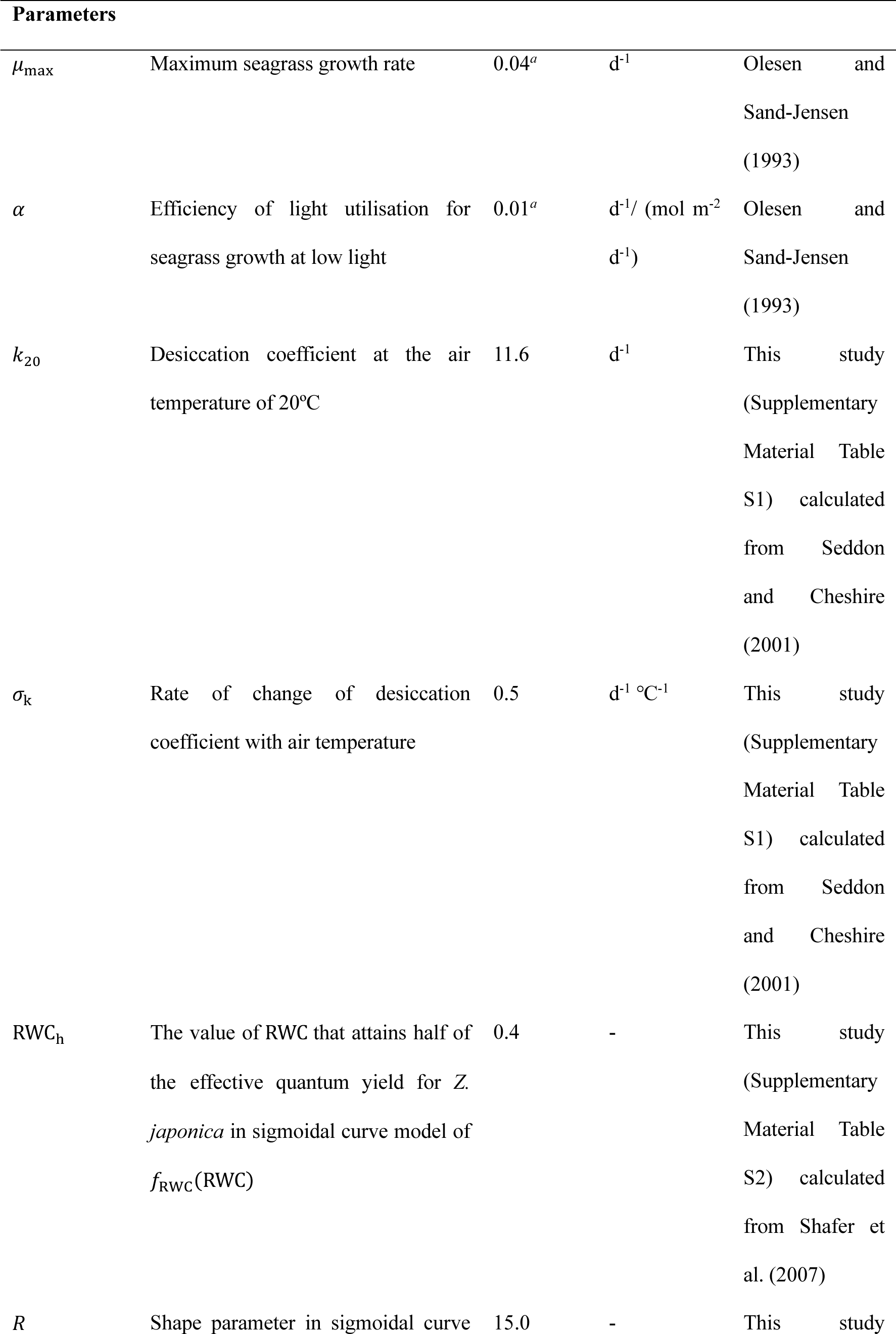
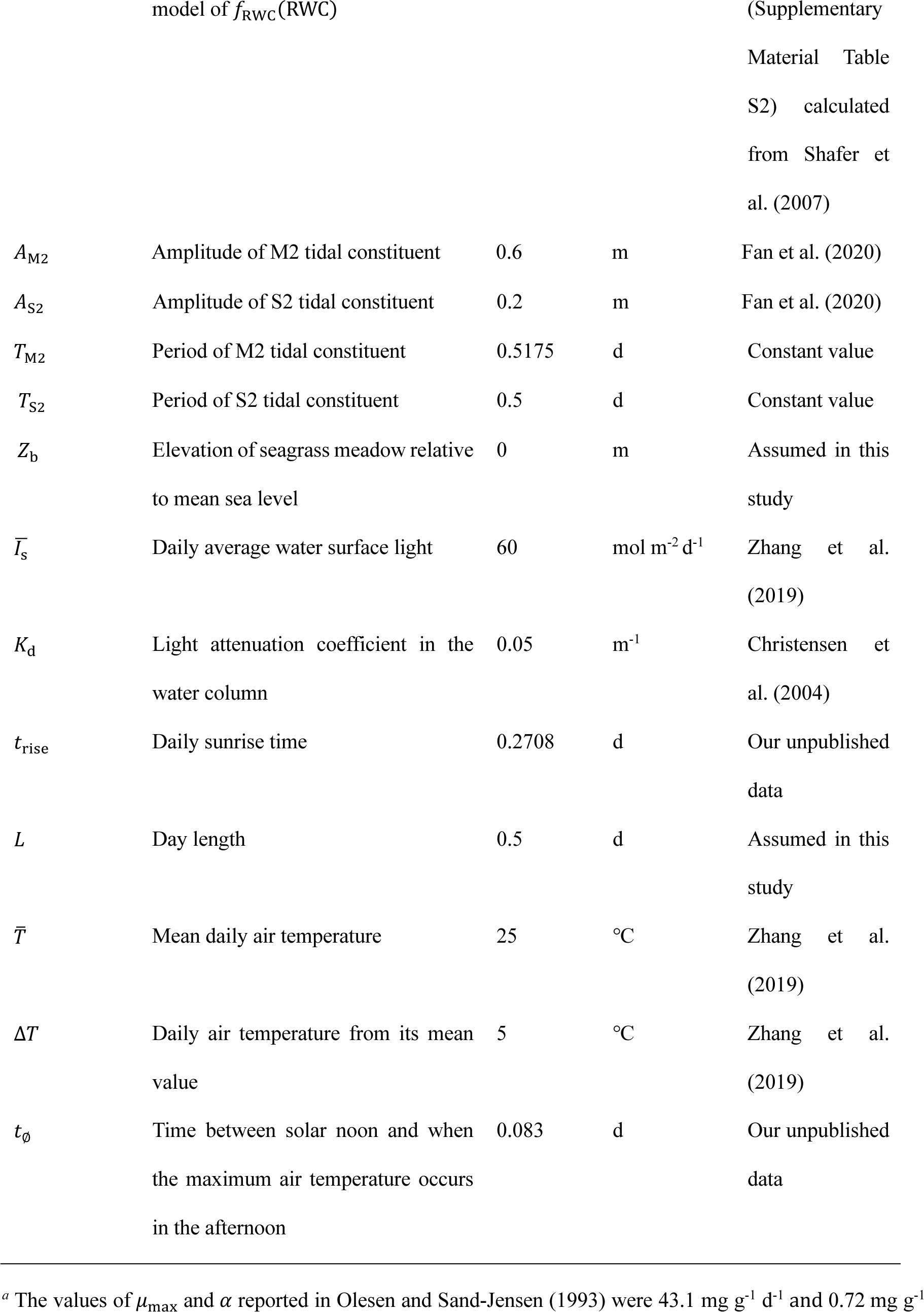
Modelling variables and parameters for the Baseline scenario.

Because we wanted to include the effects of air temperature-dependent desiccation (Eq. (8)) in our explorative simulations, and the only data available for this relationship are for two species (temperate *P. australis* and *A. antarctica*, see Supplementary Material Table S1), we parameterised air temperature-dependent desiccation (Supplementary Material Fig. S1) in the Baseline scenario based on *P. australis* (Seddon and Cheshire 2001) due to this species’ similar climatic zone to *Z. japonica*. Whilst the reported desiccation coefficient for *Z. japonica* is closer to the coefficients for *P. australis* than *A. antarctica* (Table 1), we also acknowledge that the difference in these coefficients exceeds an order of magnitude. We thus reiterate here that the purpose of our simulations is to explore consequences of the model behavior rather than parameterise the intertidal seagrass model precisely for *Z. japonica*, as air temperature-dependent desiccation coefficient information is not currently available for this species.

Finally, we chose “semi-diurnal” tides and “microtidal” conditions for the Baseline scenario, since the YRE is a microtidal estuary dominated by semi-diurnal tides (Zhou et al. 2022). Semi-diurnal tides indicate that two high tides and two low tides of similar size occur every day, and microtidal conditions indicate a daily tidal range of less than 2 m. Parameterisation of the amplitude for tidal constituents yielding these microtidal conditions was obtained from the field monitoring data in the YRE (Fan et al. 2020). The tidal periods for M2 and S2 constituents are constant. To assess the applicability of our model in the Baseline scenario, which explores the physiological responses of intertidal seagrass to desiccation while minimizing the impact of light availability, we assumed a low turbidity level (represented by water-column light attenuation coefficient *K*_d_ = 0.05 m^-1^), which is the lowest value for seagrass meadows we could find in the literature (Christensen et al. 2004). In the Baseline scenario we also assumed a meadow elevation at mean sea level, *z*_=_ = 0 m, to represent the intermediate intertidal zone. In some of our testing scenarios described later, we also examined the effects of different meadow elevations and water turbidity on intertidal seagrass growth. The full parameterisation of the Baseline scenario is given in Table 2.

**Testing scenarios.** The testing scenarios were divided into four groups (labelled as Groups I, II, III and IV, see Supplementary Material Table S3), and these scenarios were compared to the Baseline scenario. Simulations in Group I aimed to examine the effects of including air-exposure responses in seagrass growth models (i.e., *μ*(*I*) versus *μ*(*I*, RWC)), as well as the effects of different model assumptions. The model assumptions compared were the different forms of *f_RWC_*(RWC) - linear (Eq. (9)) versus hyperbolic tangent (Eq. (10)) versus sigmoidal (Eq. (11)) - and whether *μ*(*I*, RWC) follows a multiplicative formulation (Eq. (3)) or law of the minimum formulation (Eq. (4)). For testing scenarios that used the hyperbolic tangent form of *f_RWC_*(RWC), parameters for *T. hemprichii* at the air temperature of 24°C (similar climatically to YRE, Supplementary Material Table S2) were used. All testing scenarios in Group I were otherwise the same as the Baseline scenario.

Simulations in Groups II aimed to examine the effects of water turbidity (five turbidity levels from low to high) and meadow elevation (intertidal versus subtidal versus supratidal zones). More specifically, the five levels of water turbidity tested were *K*_d_ = 0.05 m^-1^, 0.5 m^-1^, 1 m^-1^, 1.5 m^-1^ and 2 m^-1^, as they represent a range of *K*_d_ values (0.05 to 2 m^-1^) corresponding to a reasonable range of irradiances that seagrasses may be able to tolerate (Christensen et al. 2004). We considered different zones along the intertidal gradient by simulating a continuous gradient of meadow elevations ranging from 3 m below mean sea level to 3 m above mean sea level (i.e., *z*_=_ ranging from −3 m to 3 m). Three different zones were subsequently categorized based on tidal range as follows: the intertidal zone spans the area between the low and high tides and is affected by daily tidal cycle; the subtidal zone is situated below the low tides and serves as a permanently submerged zone; and the supratidal zone is positioned above the high tides and is not inundated at any time. This continuous gradient of meadow elevations was simulated at the same five water turbidity levels.

Simulations in Group III were identical to simulations in Group II, except that all Group II simulations were for microtidal conditions, and all Group III simulations were for mesotidal conditions (i.e., daily tidal range between 2 m and 4 m). Similarly, Group IV simulations were identical to Group II simulations, except that Group IV simulations were performed for macrotidal conditions (i.e., daily tidal range greater than 4 m). A larger tidal range represents a wider intertidal zone and stronger tidal effects on intertidal seagrasses. The parameterisations of mesotidal and macrotidal conditions were obtained from studies on temperate intertidal Zostera meadows located in the north-western Portuguese Coast (Azevedo et al. 2017) and French Atlantic Coast (Toublanc et al. 2015), respectively (see Supplementary Material Table S3 caption for full details). Hence, the Baseline scenario and testing scenarios in Groups II, III and IV collectively provide a multifactorial simulated comparison of the effects of meadow elevation, water turbidity and tidal conditions on intertidal seagrass growth. This final targeted set of simulations allowed us to investigate (1) the trade-off between the increased light experienced further up the depth gradient (beneficial for seagrasses) and the increased desiccation further up the depth gradient (detrimental for seagrasses), and (2) how this trade-off depends on water turbidity and tidal conditions. All baseline and testing scenarios for the model application are detailed in Supplementary Material Table S3.

## 3. Results

### 3.1 The variability of seagrass photosynthetic responses to air exposures

Our first finding is that there is considerable variability in the photosynthetic responses of seagrasses to air exposures among locations and seagrass tissues. Specifically, among the same *Z. marina* species at distinct locations, substantial differences are evident in the relationship between seagrass photosynthetic efficiency and air-exposure duration, as observed in the results of experimental studies in Koje Bay and Aenggang Bay on the southern coast of South Korea (Fig. 3a). Notable distinctions also arise for *Z. japonica* when comparing our modeling results with experimental data from Padilla Bay in the USA and Koje Bay in South Korea, respectively (Fig. 3b). Furthermore, an examination of different seagrass tissue types reveals a higher desiccation tolerance in entire shoots compared to leaves for *Z. japonica*, as indicated by experimental data (Fig. 3b). The variability becomes more pronounced when comparing our modeled results and experimental data for *Z. marina*, considering both different tissue types and locations (Fig. 3a). This suggests that it is difficult to transfer a model parameterised at one location/for one seagrass tissue to another location/tissue due to the influence of environmental conditions such as light, temperature, wind, humidity, soil properties, etc. in the field as well as the different water retention ability of seagrass tissues (Kim et al. 2020; Suykerbuyk et al. 2018).

**Fig. 3.**
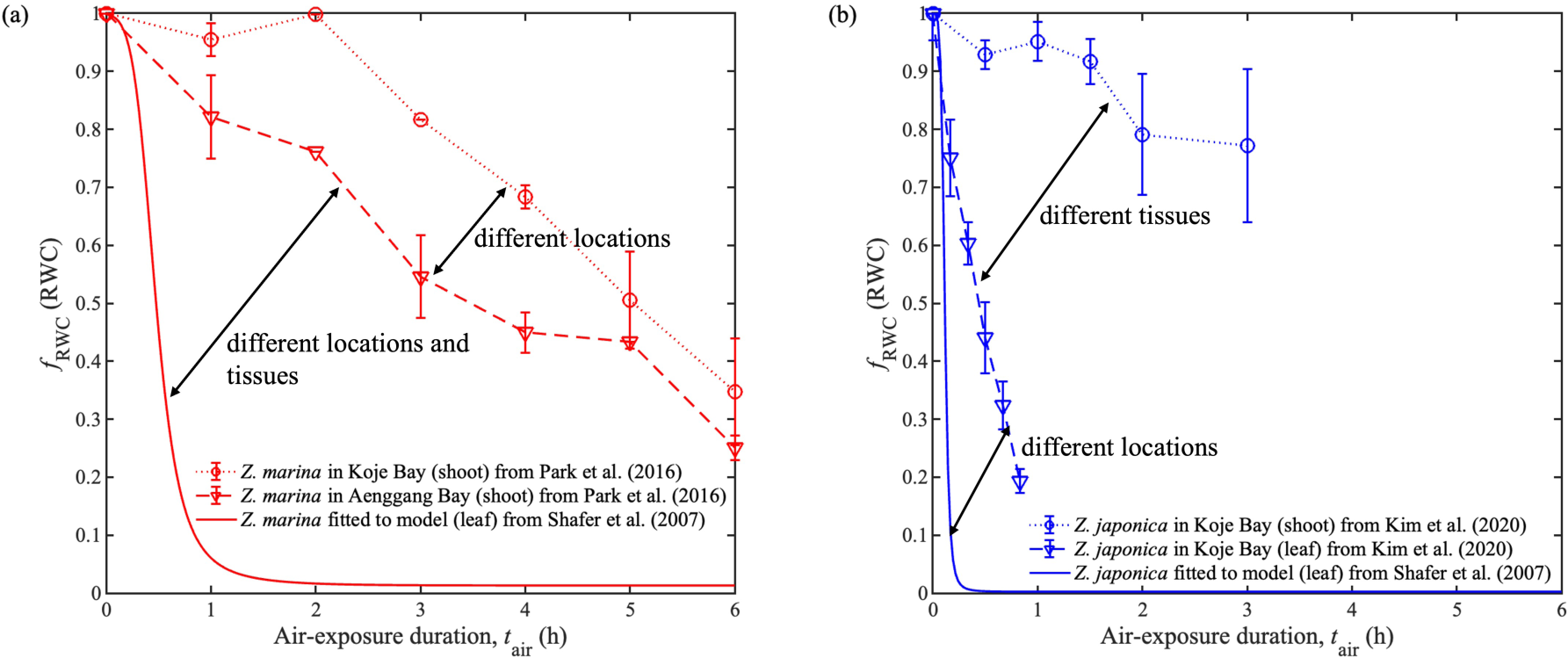
Relationship between seagrass photosynthetic efficiency and air-exposure duration for (a) *Z. marina* and (b) *Z. japonica.* Notice that there are substantial differences in RWC versus air-exposure duration for the same species at different locations and in different tissues (shoot vs leaf).

### 3.2 Physiological responses of intertidal seagrasses when air-exposed

Our finding is that desiccation has a substantial effect on seagrass growth rate, thus justifying the new formulations for growth rate *μ*(*I*, RWC) introduced in the present work, as follows. In our Baseline scenario of simulating intertidal seagrasses (Eqs. (6) - (11), (12) - (16) parameterized using Table 2), the modelled water depth fluctuates throughout the spring-neap tidal cycle (Fig. 4a). The intertidal seagrasses are subject to air exposure at low tides twice a day, each lasting for nearly 6 hours (Supplementary Material Fig. S9a), with the RWC dropping to low values in each exposure period (Supplementary Material Fig. S9b). As a result, the seagrass growth rate in the Baseline scenario, which is dependent on both light and RWC (*μ*(*I*, RWC)), fluctuates throughout the spring-neap tides (Fig. 4b). On the other hand, if the seagrass growth rate depends solely on light (*μ*(*I*)), the fluctuation of this growth rate due to tidal-induced light deprivation is very minor (Fig. 4c). Furthermore, the modelled average growth rate (represented by the dashed lines in Fig.4) is substantially lower for *μ*(*I*, RWC) than for *μ*(*I*).

**Fig. 4.**
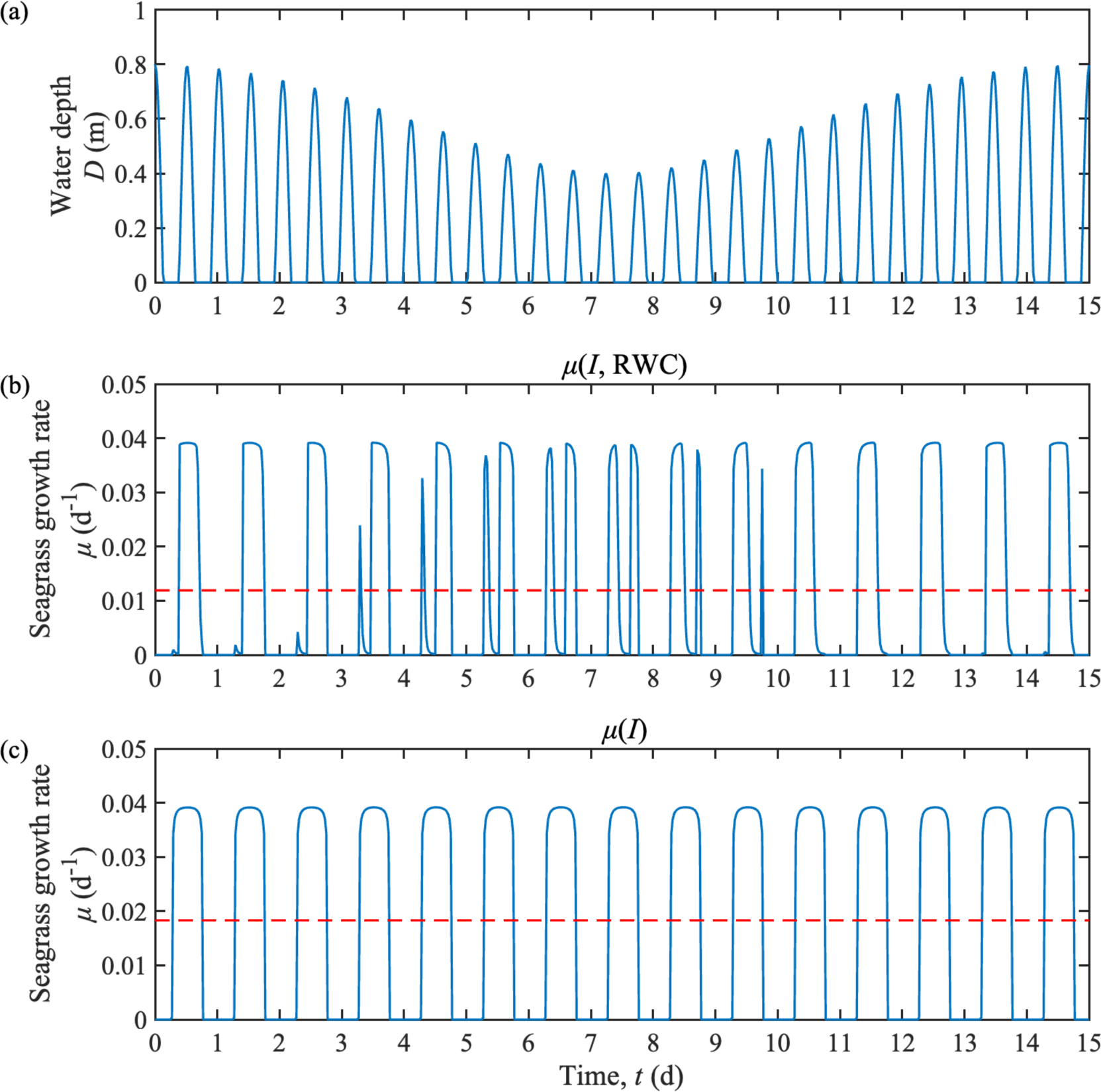
(a) Modelled water depth over 15 d in the Baseline scenario (Table 2); Predicted seagrass growth rate over 15 d for (b) growth rate dependent on light and RWC (Baseline scenario, see Table 2) and (c) growth rate only dependent on light (first testing scenario in Group I, see Supplementary Material Table S3). The red dashed line represents the mean value of the growth rate over the 15-d simulation. Notice that accounting for desiccation causes the prediction of the mean growth rate to be substantially lower (compare red dashed lines between (a) and (b)).

Next, we investigated how the cumulative effect of air-exposure duration and light deprivation on seagrass growth rate is mediated by the multiplicative or law of the minimum formulation representing these cumulative stressors. To accomplish this, we performed simulations that exactly matched the Baseline scenario (Table 2) or possessed minor modifications of this scenario. More specifically, for a single exposure-inundation cycle, we adopted two constant values of benthic light irradiance in the model, i.e. one below and one above the saturating irradiance (15 mol m^-2^ d^-1^ in Park et al. (2021)), and we selected a single exposure-inundation cycle during spring tide in our model scenarios (Fig. 5).

**Fig. 5.**
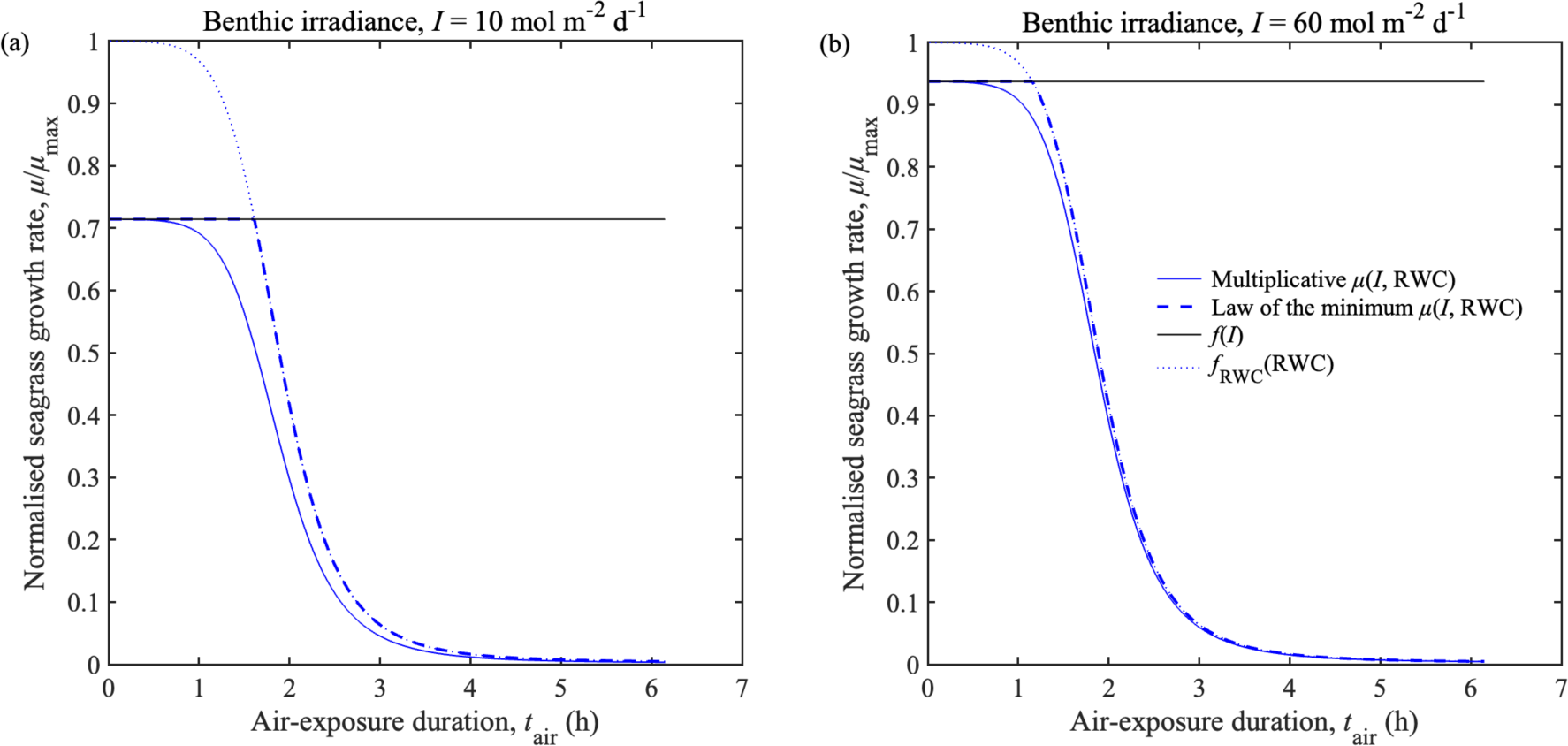
The relationship between the normalized growth rate (*μ*/*μ*_max_) and air-exposure duration (*t*_air_) for two seagrass growth functions with sigmoidal curve *f_s_*_;<=_(RWC) under two light irradiance conditions: (a) 10 mol m^-2^ d^-1^ and (b) 60 mol m^-2^ d^-1^. These simulations are equivalent to, or slight modifications of the Baseline scenario described in Table 2. Notice that the growth rate always reduces with air-exposure duration faster with the multiplicative formulation (solid blue line) is assumed, compared to the law of the minimum formulation (dashed blue line), but this effect is reduced as benthic irradiance increases (e.g., from panel (a) to (b)).

We utilized the normalized growth rate (*μ*/*μ*_$%&_) to compare the differences between growth rate scenarios using the multiplicative or law of the minimum formulation for *Zostera* spp. with sigmoidal *f_s_*_()*_(RWC). The results (Fig. 5) reveal that, regardless of the light level, when applying the multiplicative formulation (solid blue lines), the growth rate begins to decline earlier in response to air exposure than when applying the law of the minimum formulation (dashed blue lines). However, when the benthic light irradiance is 60 mol m^-2^ d^-1^ (well above the saturating irradiance, Fig. 5b), the differences in growth rate response to air exposure between multiplicative and law of the minimum formulation scenarios are smaller than when the benthic irradiance is below saturation (10 mol m^-2^ d^-1^, Fig. 5a). The law of the minimum formulation consistently yields a more optimistic prediction of seagrass growth rate than the multiplicative formulation, and the difference between the predictions of these two formulations tends to widen when multiple stressors (e.g., desiccation and light deprivation) are expected to have substantial detrimental impacts on seagrass growth.

### 3.3 Species-specific effects on intertidal seagrass growth

We next ran simulations with three different functional forms *f_RWC_*(RWC) for the effect of seagrass tissue relative water content on seagrass photosynthetic efficiency to examine the impact of desiccation tolerance of different seagrass species on their growth rates (Baseline and Group I scenarios, see Supplementary Material Table S3). The results show that the hyperbolic tangent model assumed for *f_RWC_*(RWC) yielded a higher predicted 15d-averaged growth rate of seagrass in comparison to other functional forms of *f_RWC_*(RWC) (Fig. 6). This suggests that the function *f_RWC_*(RWC) characterizing the desiccation tolerance of species constitutes an important factor. The reason that the hyperbolic tangent model for *f_RWC_*(RWC) yields a higher overall growth rate for intertidal seagrass is because mathematically it predicts a higher photosynthetic efficiency *f_RWC_*(RWC) at all values of RWC < 1 compared to the other two functional forms (linear and sigmoidal curve). The difference in *f_RWC_*(RWC) between the hyperbolic tangent model and the other two forms are particularly large at high RWC values which are experienced by the seagrasses soon after air exposure begins. Supplementary Material Fig. S11 demonstrates that this causes a delay in the reduction of seagrass growth rate following air exposure if the hyperbolic tangent model of *f_RWC_*(RWC) is assumed (green lines in Supplementary Material Fig. S11); this delay in the reduction of seagrass growth rate following air exposure is substantially reduced for the other two functional forms of *f_RWC_*(RWC) (blue and red lines in Supplementary Material Fig. S11). Each *f_RWC_*(RWC) was obtained from different seagrass species (Supplementary Material Fig. S5-S7), and thus it may be the case that species-specific functional forms of *f_RWC_*(RWC) play a key role in determining the seagrass species tolerance to desiccation.

**Fig. 6.**
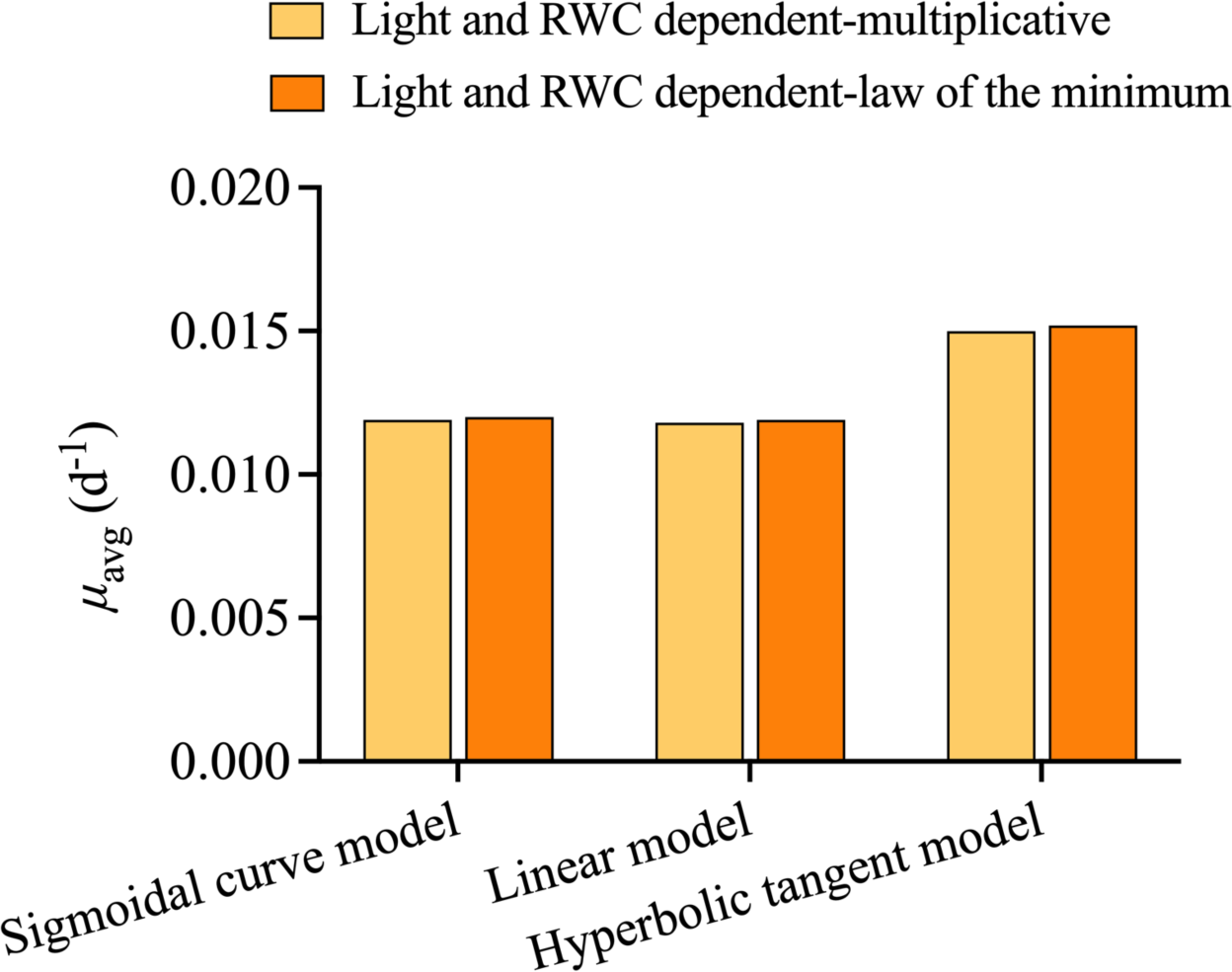
The results of modelled average growth rate (*μ*_max_) during the simulation period of 15 d for light and RWC dependent growth rate followed the multiplicative or law of the minimum formulation with the three types of *f_s_*_;<=_(RWC) defined in Eq. (9)-(11). Notice that mean growth rates are always predicted to be higher if the hyperbolic tangent form of *f_s_*_;<=_(RWC) (i.e., Eq. (10) is assumed.

Meanwhile, there were no substantial differences between average growth rates for functions that followed the multiplicative or law of the minimum formulations with the same *f_RWC_*(RWC). Note that these growth rates were calculated under very high light conditions (60 mol m^-2^ d^-1^as assumed in the Baseline scenario), which led to this small difference between the multiplicative and law of the minimum formulations (see Fig. 5b and the related discussion). At lower irradiances (10 mol m^-2^ d^-1^), these differences increase marginally (see Supplementary Material Fig. S10), and the difference between functional forms of *f_RWC_*(RWC) tends to have a larger effect on the growth rate (Supplementary Material Fig. S11a).

### 3.4 Intertidal seagrass growth along vertical gradient under different tidal range condition

Within a 15-d simulation in different modelling scenarios (Baseline and Groups II, III and IV, see Section 2.4.2 and Supplementary Material Table S3 for full details), we compared the 15d-averaged predictions of (1-*f_s_*(*I*)) and (1-*f_RWC_*(RWC)) to illustrate, and as metrics of, the light deprivation and desiccation stress on the intertidal seagrass growth, respectively. The results show that within a specific tidal range area in the intertidal zones, light deprivation stress gradually decreases and desiccation stress increases as meadow elevation increases (compare red and blue lines in Fig.7 a, c & e). Meanwhile, the light deprivation stress on seagrass growth dramatically increases as the water turbidity level rises (compare different red lines in Fig. 7a, c & e), leading to more substantial variations along the vertical gradient. In the subtidal zones, light deprivation is the predominant stress on seagrass growth, which gradually decreases as elevation increases, and desiccation stress is absent in these zones. In contrast, desiccation is the dominant stressor impacting seagrass growth in the supratidal zones, where light deprivation stress is minimal (only limited by the efficiency of light utilisation for seagrass growth).

**Fig. 7.**
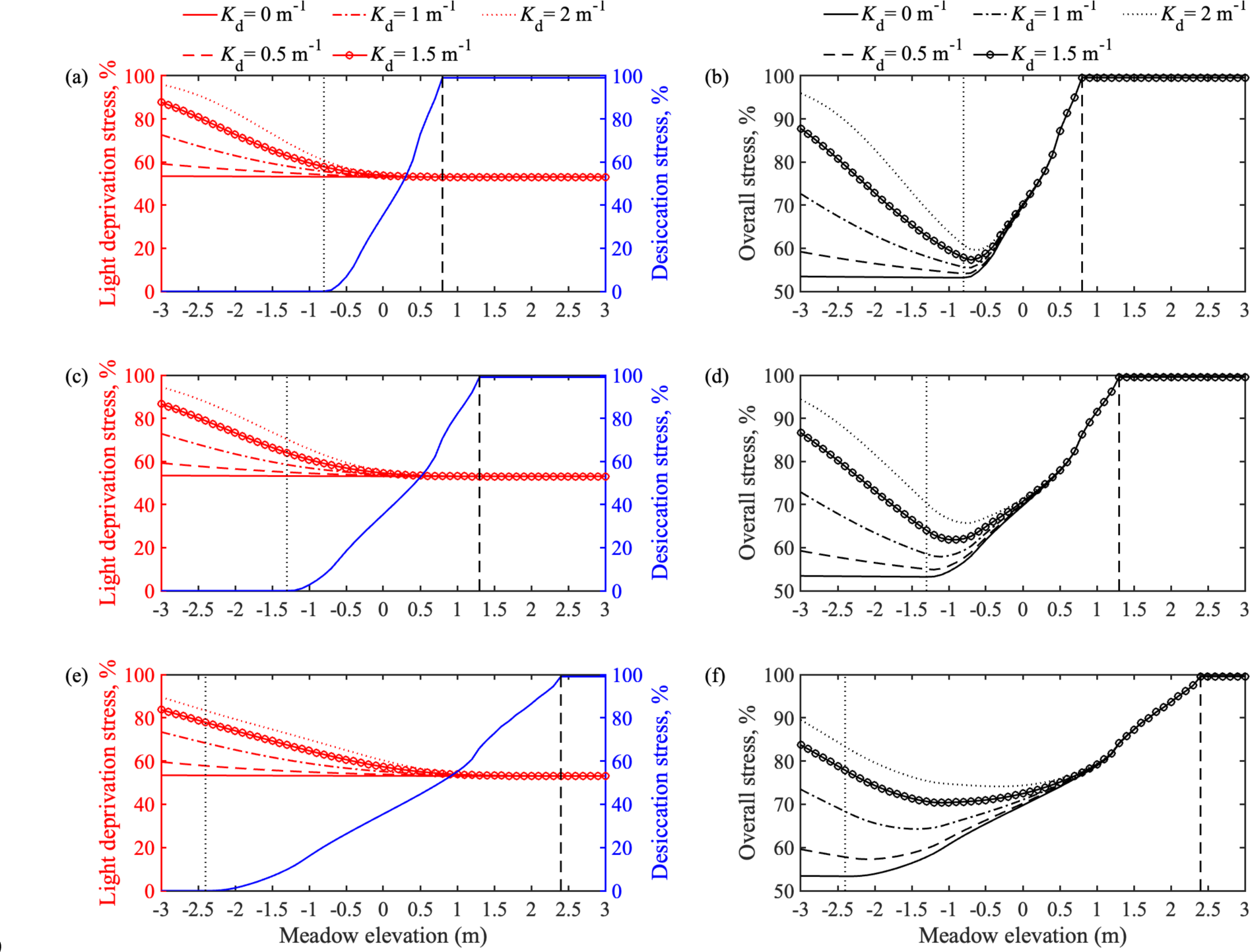
The modelled results of light deprivation stress and desiccation stress on 15 d-averaged seagrass growth with multiplicative formulation along the vertical depth gradient under different water turbidity conditions and tidal conditions. The separate effects of light deprivation stress and desiccation stress are shown in panels (a, c, e) and the combined effect of both stressors is shown in panels (b, d, f). Predictions are shown for microtidal conditions (a, b), mesotidal conditions (c, d), and macrotidal conditions (e, f). In each of the six panels, the black dotted line is the boundary between subtidal and lower intertidal zones, while the black dashed line is the boundary between upper intertidal and supratidal zones. The intertidal zone extends from −0.8 m to 0.8 m, −1.3 m to 1.3 m, and −2.4 m to 2.4 m for micro-, meso- and macro-tidal areas, respectively; The subtidal zone extends below −0.8 m, −1.3 m, and −2.4 m for micro-, meso- and macro-tidal areas, respectively; The supratidal zone extends above 0.8 m, 1.3 m, and 2.4 m for micro-, meso- and macro-tidal areas, respectively. Meadow elevation *z*_b_(m) is relative to mean sea level, and *K*_d_ (m^-1^) is the light attenuation coefficient of the water column. These plots were constructed using the Baseline and Groups II, III and IV modelling scenarios (Supplementary Material Table S3) that assumed the multiplicative formulation for *μ*(*I*, RWC).

In addition, the overall stress on intertidal seagrass growth, resulting from the cumulative effects of light deprivation and desiccation, was simulated using the reduction in seagrass growth rate from its potential maximum value, i.e., (1-*μ*/*μ*_max_) as a metric representing this overall stress (Fig. 7b, d & f). The results indicate the presence of an “optimal” elevation for intertidal seagrasses where the cumulative stress arising from desiccation and light deprivation is minimized (e.g., at a meadow elevation approximately 0.5 m below mean sea level for seagrasses growing in microtidal conditions with water turbidity of *K*_d_ = 2 m^-1^ as shown in Fig. 7b). At meadow elevations below the predicted optimal elevation, the overall stress increases with turbidity. This suggests that below the optimal elevation, the growth of intertidal seagrasses is primarily limited by light availability, and air exposure acts as the ‘window’ of photosynthetic relief from high turbidity to mediate the light deprivation stress. As elevation of the seagrass increases above the optimal elevation, the difference in overall stress between the different turbidity scenarios gradually diminishes. Thus, for seagrasses growing at elevations above the optimal elevation, desiccation becomes the dominant limiting factor for growth, and the negative effects of air exposure increasingly outweigh the positive effects of increased light moving further upward. Across all model scenarios we tested, the predicted optimal elevation shifts to lower elevations (more specifically, towards the lower intertidal zones), as turbidity reduces, although the curves tend to flatten. Note also that using the law of the minimum formulation does not change the conclusion (Supplementary Material Fig. S12).

The results confirm that light is a primary control of intertidal seagrass growth in the subtidal zones, while desiccation is a primary control in the supratidal zones. Our findings clearly suggest a trade-off between light deprivation and desiccation in relation to intertidal seagrass growth along the intertidal gradient, with an optimal elevation for seagrasses situated within the intertidal zone that maximises the benefits of light availability whilst minimising the detrimental effects of desiccation (Fig. 7b, d & f).

## 4. Discussion

### 4.1 Species-specific physiological responses of intertidal seagrasses when air-exposed

Previous experimental studies have demonstrated the significance of physiological processes such as RWC loss and associated photosynthesis decline of different intertidal seagrass species when subject to air exposure (Jiang et al. 2014; Shafer et al. 2007). However, it is still uncertain how these physiological processes further limit the growth rate of intertidal seagrasses under different environmental conditions such as tidal range, meadow elevation and water turbidity. Our study introduced new data-calibrated formulations for intertidal seagrasses that quantify the decline of photosynthetic efficiency due to the changes of RWC when air-exposed. To do this, a comprehensive review of the literature on seagrass desiccation (Table 1) and the effects of this desiccation on photosynthetic efficiency (Supplementary Material Fig. S5-S7) was conducted. We then undertook targeted model simulations which demonstrated that air-exposed physiological processes (light- and RWC-dependent growth) can be substantially lower than if the air-exposed responses are neglected (light-dependent growth) (dashed lines in Fig. 4). This suggests that neglecting the physiological response to air exposure can yield overestimation of growth rates for intertidal seagrasses.

We also examined the species-specific effect of desiccation on the growth rate of intertidal seagrasses by comparing the growth rate predictions made using three different *f_s_*_()*_(RWC) functions (Eq. (9)-(11)) characterizing the changes of photosynthetic efficiency with the temporal decline of RWC due to short-term air exposure. These three *f_s_*_()*_(RWC) functions are data-calibrated functions that we collate, justify and introduce in the present work. Our results suggest that the choice of the function *f_s_*_()*_(RWC) characterizing the desiccation tolerance of different species substantially affects quantitative predictions of the physiological responses of intertidal seagrasses when air-exposed; if these functions are indeed species-specific, they may also suggest which species are more or less tolerant to desiccation. For example, desiccation-sensitive seagrass species such as temperate *H. johnsonii* and *C. nodosa* (Kahn and Durako 2009; Papathanasiou et al. 2020) as well as temperate *Z. japonica* and *Z. marina* (Shafer et al. 2007), whose response in photosynthetic efficiency is more likely to follow a linear (Eq. (9)) or sigmoidal curve (Eq. (11)) in relation to RWC loss, would experience an immediate and rapid decrease in growth rate following air exposure (red and blue lines in Supplementary Material Fig. S11). In contrast, desiccation-tolerant species such as tropical *T. hemprichii* and *E. acoroides* (Jiang et al. 2014), whose response in photosynthetic efficiency is more likely to follow a hyperbolic tangent (Eq. (10)) in relation to RWC loss, may tolerate hours of air exposure without affecting their growth (green lines in Supplementary Material Fig. S11). However, we could only find data for six species to parameterise these *f_s_*_()*_(RWC) functions. In the future, more species-specific studies can be incorporated; for example Bjork et al. (1999) reported that tropical intertidal seagrass species were more desiccation-resistant and were likely to have higher tolerances to thrive in the intertidal zones.

### 4.2 Trade-off between light deprivation and desiccation related to intertidal seagrass distribution

Along the intertidal gradient, both light availability and stress of desiccation gradually increase as meadow elevation increases. Light is a primary control on seagrass growth in subtidal zones, while desiccation is a primary control on seagrass growth in supratidal zones. However, in the intertidal zones, we observed in our simulations a clear trade-off between light deprivation and desiccation along the intertidal gradient (Fig. 7), as a balance between obtaining enough light for their growth, while also avoiding the detrimental consequences of desiccation stress is vital for intertidal seagrasses. This is thus an “optimal” elevation for intertidal seagrasses which represents the minimized combined stress of light deprivation and desiccation. The location of this optimal elevation for intertidal seagrasses varies under different environmental conditions, such as tidal range and water turbidity. In all our model scenarios, the predicted optimal elevation occurs in the lower or intermediate intertidal zones. As water turbidity increases, the optimal elevation shifts upwards to higher elevations. The evaluation of optimal elevation has the potential to inform the most suitable habitat for intertidal seagrass growth.

Previous studies have found that differences in desiccation tolerances can be responsible for the seagrass distribution along the intertidal gradient. For example, in the intertidal seagrass meadows at the coasts of the Indo-Pacific, the desiccation-tolerant species *T. hemprichii* was found to be dominant in the upper intertidal zone while desiccation-sensitive species *H. uninervis* occupied the lower intertidal zone (Lan et al. 2005). However, increasing evidence suggests that photosynthetic responses to desiccation are insufficient to explain observed patterns of intertidal zonation (Shafer et al. 2007). Therefore, it is necessary to consider additional mechanisms, such as the combined effect of desiccation and light deprivation considered here, to explain the observed zonation patterns of intertidal seagrasses. The trade-off between light deprivation and desiccation on intertidal seagrass distribution also finds some agreement with field studies demonstrating the intertidal zonation of different seagrass species. Huong et al. (2003) found that intertidal *Z. japonica* in northern Vietnam occupied the intermediate intertidal zone while *H. ovalis* dominated in the lower intertidal zone, due to the different tolerances to low light availability (less in *Z. japonica*) and desiccation (less in *H. ovalis*). Meanwhile, the seagrass *Z. japonica* was also found to have the highest biomass in the intermediate intertidal zone on the southern coast of South Korea, where air exposure and light availability determined the upper and lower distributional limits, respectively (Kim et al. 2016). However, other environmental factors may also play an important role in defining the zonation of seagrass colonisation; Infantes et al. (2009) suggests that subtidal seagrasses have an upper depth limit controlled by shallow-water wave action, but the relevance of this limit to intertidal seagrasses may depend on the harshness of the local hydrodynamic conditions. Notably, intertidal seagrasses also evolve adaptation mechanisms to air exposure stress through adjustments to physiological characteristics followed by changes to morphology (Manassa et al. 2017; Park et al. 2016). For example, the enhanced photosynthetic performance after air exposure and the layout of the densely overlapped leaves to attain water are attributed as the adaptation mechanisms for *Z. japonica* in the intertidal zone (Kim et al. 2020). Regardless of these complexities, understanding the trade-offs between stressors influencing the lower and upper meadow elevations of seagrasses is crucial for the effective management of these habitats, especially since intertidal zones are dynamic and challenging environments.

### 4.3 Model applications and future work

Our study emphasizes the importance of understanding the air-exposed physiological responses on the growth dynamics of intertidal seagrasses, and the growth rate functions we introduce can be immediately incorporated into a wide variety of process-based seagrass growth models. Although we are here only considering the impact of light and RWC on seagrass growth, the future application of the model components we introduce could also incorporate other interacting factors (e.g., temperature, nutrients) by the inclusion of appropriately defined additional functions (Baird et al. 2016; Elkalay et al. 2003; Turschwell et al. 2022). Our study provides conceptual and mathematical guidance for ecological modellers to include air-exposed responses of intertidal seagrasses in their coastal ecosystem models. One example future application of our intertidal seagrass growth dynamics, of substantial interest, could be to simulate scenarios that assist in the selection of suitable sites for seagrass transplanting. Additionally, in recent years, global sea level rise and an increase in the input of terrestrial sediments pose a hazard to intertidal seagrass ecosystems (Flowers et al. 2023). These stressors simultaneously change the tidal regime and water turbidity, which affects the duration of air exposure/inundation periods and light availability. The model formulations we discuss in the present work can account for these cumulative stressors.

The comparisons between our data-calibrated model results and additional experimental studies also demonstrated the variability of physiological responses of intertidal seagrasses to air exposure, and hence the difficulty in transferring a model parameterised at one location/for one seagrass tissue to another location/tissue (Fig. 3). This suggests that future experimental studies on the relationship between photosynthetic efficiency and air-exposure duration may therefore need to be species-or location-specific, to improve model predictions, although we recognise that this may often be prohibitively difficult or expansive to implement.

Future modelling studies for intertidal seagrasses could also further incorporate delayed recovery processes of photosynthetic efficiency after re-submersion. Recovery of photosynthetic efficiency after re-submersion is critical for the seagrass growth response to desiccation stress (Park et al. 2016; Seddon and Cheshire 2001; Shafer et al. 2007). When intertidal seagrasses are exposed to air for a prolonged duration, their photosynthetic efficiency may not be able to recover to their initial level after re-submersion. In the worst-case scenario, seagrasses may even lose the ability to resume photosynthesis (Shafer et al. 2007). There is not yet sufficient quantitative information available in the literature for us to confidently propose models of recovery of photosynthetic efficiency after re-submersion; this is an open question for future experimental research.

## 5. Conclusion

Through a comprehensive review of seagrass desiccation literature and the subsequent development of the first (to our knowledge) formulation of seagrass growth responses to air exposure, our study was able to explore how seagrass growth dynamics is affected by periodic tidal inundation and exposure under a wide range of environmental scenarios (tidal range, meadow elevation and water turbidity). We showed that neglecting physiological responses to air exposure for intertidal seagrasses results in overestimated growth rates, and we revealed a trade-off between light deprivation and desiccation on seagrass growth along intertidal gradients. More specifically, there is an “optimal” elevation for seagrasses where the combined stressors of desiccation and light deprivation are minimized, although the precise location of this optimal elevation will be highly system-specific. This finding may have future application in evaluating the viability of intertidal seagrass habitats and in informing site selection for seagrass transplanting. Overall, our work highlights the importance of elucidating the physiological responses of intertidal seagrasses in a highly dynamic and harsh environment and prompts further experimental studies to inform improved modeling of intertidal seagrass growth.

## Acknowledgments

This work was supported by the National Key R&D Program of China (2019YFE0121500), the Australian Research Council (ARC) Discovery Early Career Researcher Award DE200100683, and the Scholarship from China Scholarship Council. The authors thank Severine Choukroun, Catherine Collier and Lucas Langlois for helpful discussions during manuscript development.

## Data Availability

The main code supporting the model simulation of this article is available online at https://github.com/ceeh-bnu/Intertidal-seagrass.

## Author Contribution Statement

X.W. contributed to conceptualization, model design, data analysis and interpretation, and drafting and revising manuscript. M.A and D.S. contributed to conceptualization, data interpretation, and reviewing and editing manuscript. All authors contributed to and accepted the final version of the manuscript.

## References

Adams, J., and G. Bate. 1994. The tolerance to desiccation of the submerged macrophytes Ruppia cirrhosa (Petagna) Grande and Zostera capensis Setchell. J. Exp. Mar. Biol. Ecol. 183: 53–62. doi: 10.1016/0022-0981(94)90156-2

Adams, M. P. and others. 2020. Predicting seagrass decline due to cumulative stressors. Environ. Model. Softw. 130: 104717. doi: 10.1016/j.envsoft.2020.104717

Azevedo, A., A. I. Lillebø, J. Lencart e Silva, and J. M. Dias. 2017. Intertidal seagrass models: Insights towards the development and implementation of a desiccation module. Ecol. Model. 354: 20–25. doi: 10.1016/j.ecolmodel.2017.03.004

Baird, M. E. and others. 2016. A biophysical representation of seagrass growth for application in a complex shallow-water biogeochemical model. Ecol. Model. 325: 13–27. doi: 10.1016/j.ecolmodel.2015.12.011

Balke, T., M. Stock, K. Jensen, T. J. Bouma, and M. Kleyer. 2016. A global analysis of the seaward salt marsh extent: The importance of tidal range. Water Resour. Res. 52: 3775–3786. doi: 10.1002/2015wr018318

Bertelli, C. M., and R. K. F. Unsworth. 2018. Light Stress Responses by the Eelgrass, Zostera marina (L). Front. Environ. Sci. 6: 39. doi: 10.3389/fenvs.2018.00039

Bjork, M., J. Uku, A. Weil, and S. Beer. 1999. Photosynthetic tolerances to desiccation of tropical intertidal seagrasses. Mar. Ecol. Prog. Ser. 191: 121–126. doi: 10.3354/meps191121

Cabaço, S., R. Machás, and R. Santos. 2009. Individual and population plasticity of the seagrass Zostera noltii along a vertical intertidal gradient. Estuarine, Coastal Shelf Sci. 82: 301–308. doi: 10.1016j.ecss.2009.01.020

Carr, J. A., P. D’Odorico, K. J. McGlathery, and P. L. Wiberg. 2012. Stability and resilience of seagrass meadows to seasonal and interannual dynamics and environmental stress. J. Geophys. Res.: Biogeosci. 117: 1007. doi: 10.1029/2011JG001744

Che, X., H. Li, L. Zhang, and J. Liu. 2022. Effect of high light and desiccation on photosystem II in the seedlings and mature plants of tropical seagrass Enhalus acoroides during low tide. J. Oceanol. Limnol. 41: 241–250. doi: 10.1007s00343-021-1170-2

Christensen, P., E. Díaz Almela, and O. Diekmann. 2004. Can transplanting accelerate the recovery of seagrasses? J. Borum, C. Duarte, D. Krause-Jensen, T.M. Greve (Eds.), European Seagrasses: an Introduction to Monitoring and Management, M&MS Project, Copenhagen, pp. 77-82

Clavier, J. and others. 2011. Aerial and underwater carbon metabolism of a Zostera noltii seagrass bed in the Banc d’Arguin, Mauritania. Aquat. Bot. 95: 24–30. doi: 10.1016/j.aquabot.2011.03.005

Colomer, J., and T. Serra. 2021. The World of Edges in Submerged Vegetated Marine Canopies: From Patch to Canopy Scale. Water. 13: 2430. doi: 10.3390/w13172430

de los Santos, C. B. and others. 2022. Sedimentary Organic Carbon and Nitrogen Sequestration Across a Vertical Gradient on a Temperate Wetland Seascape Including Salt Marshes, Seagrass Meadows and Rhizophytic Macroalgae Beds. Ecosystems. 26: 826–842. doi: 10.1007/s10021-022-00801-5

Elkalay, K. and others. 2003. A model of the seasonal dynamics of biomass and production of the seagrass Posidonia oceanica in the Bay of Calvi (Northwestern Mediterranean). Ecol. Model. 167: 1–18. doi: 10.1016S0304-3800(03)00074-7

Erftemeijer, P. L. A., J. van Gils, M. B. Fernandes, R. Daly, L. van der Heijden, and P. M. J. Herman. 2023. Habitat suitability modelling to improve understanding of seagrass loss and recovery and to guide decisions in relation to coastal discharge. Mar. Pollut. Bull. 186: 114370. doi: 10.1016/j.marpolbul.2022.114370

Espadero, A. D. A., Y. Nakamura, W. H. Uy, P. Tongnunui, and M. Horinouchi. 2020. Tropical intertidal seagrass beds: An overlooked foraging habitat for fishes revealed by underwater videos. J. Exp. Mar. Biol. Ecol. 526: 151353. doi: 10.1016/j.jembe.2020.151353

Fan, Y., S. Chen, S. Pan, and S. Dou. 2020. Storm-induced hydrodynamic changes and seabed erosion in the littoral area of Yellow River Delta: A model-guided mechanism study. Cont. Shelf Res. 205. doi: 10.1016/j.csr.2020.104171

Flowers, G. J. L., H. R. Needham, R. H. Bulmer, A. M. Lohrer, and C. A. Pilditch. 2023. Going under: The implications of sea-level rise and reduced light availability intertidal primary production. Limnol. Oceanogr. 68: 1301–1315. doi: 10.1002/lno.12347

Folmer, E. O. and others. 2012. Seagrass–Sediment Feedback: An Exploration Using a Non-recursive Structural Equation Model. Ecosystems. 15: 1380–1393. doi: 10.1007/s10021-012-9591-6

Geary, W. L. and others. 2020. A guide to ecosystem models and their environmental applications. *Nat*. Ecol. Evol. 4: 1459–1471. doi: 10.1038/s41559-020-01298-8

Huong, T. T. L. and others. 2003. Seasonality and depth zonation of intertidal Halophila ovalis and Zostera japonica in Ha Long Bay (northern Vietnam). Aquat. Bot. 75: 147–157. doi: 10.1016S0304-3770(02)00172-9

Infantes, E., J. Terrados, A. Orfila, B. Cañellas, and A. Álvarez-Ellacuria. 2009. Wave energy and the upper depth limit distribution of Posidonia oceanica. Bot. Mar. 52: 419–427. doi: 10.1515/BOT.2009.050

Jarvis, J. C., M. J. Brush, and K. A. Moore. 2014. Modeling loss and recovery of Zostera marina beds in the Chesapeake Bay: The role of seedlings and seed-bank viability. Aquat. Bot. 113: 32–45. doi: 10.1016j.aquabot.2013.10.010

Jassby, A. D., and T. Platt. 1976. Mathematical formulation of the relationship between photosynthesis and light for phytoplankton. Limnol. Oceanogr. 21: 540–547. doi: 10.4319/lo.1976.21.4.0540

Jiang, Z., X. Huang, J. Zhang, C. Zhou, Z. Lian, and Z. Ni. 2014. The effects of air exposure on the desiccation rate and photosynthetic activity of Thalassia hemprichii and Enhalus acoroides. Mar. Biol. 161: 1051–1061. doi: 10.1007/s00227-014-2398-6

Johnson, I. R., and J. H. M. Thornley. 1984. A model of instantaneous and daily canopy photosynthesis. J. Theor. Biol. 107: 531–545. doi: 10.1016/S0022-5193(84)80131-9

Kahn, A. E., and M. J. Durako. 2009. Photosynthetic tolerances to desiccation of the co-occurring seagrasses Halophila johnsonii and Halophila decipiens. Aquat. Bot. 90: 195–198. doi: 10.1016/j.aquabot.2008.07.003

Kaldy, J. 2012. Influence of light, temperature and salinity on dissolved organic carbon exudation rates in Zostera marina L. Aquat Biosyst. 8: 19. doi: 10.1186/2046-9063-8-19

Kim, J.-H., S. H. Kim, Y. K. Kim, J.-I. Park, and K.-S. Lee. 2016. Growth dynamics of the seagrass Zostera japonica at its upper and lower distributional limits in the intertidal zone. *Estuarine*, Coastal Shelf Sci. 175: 1–9. doi: 10.1016/j.ecss.2016.03.023

Kim, S. H., J. W. Kim, Y. K. Kim, S. R. Park, and K. S. Lee. 2020. Factors controlling the vertical zonation of the intertidal seagrass, Zostera japonica in its native range in the northwestern Pacific. Mar. Environ. Res. 157: 104959. doi: 10.1016/j.marenvres.2020.104959

Kirk, J. T. 1985. Effects of suspensoids (turbidity) on penetration of solar radiation in aquatic ecosystems. Hydrobiologia. 125: 195–208. doi: 10.1007/BF00045935

Koch, E. W. 2001. Beyond light: physical, geological, and geochemical parameters as possible submersed aquatic vegetation habitat requirements. Estuaries. 24: 1–17. doi: 10.2307/1352808

Lan, C. Y., W. Y. Kao, H. J. Lin, and K. T. Shao. 2005. Measurement of chlorophyll fluorescence reveals mechanisms for habitat niche separation of the intertidal seagrasses Thalassia hemprichii and Halodule uninervis. Mar. Biol. 148: 25–34. doi: 10.1007/s00227-005-0053-y

Leuschner, C., S. Landwehr, and U. Mehlig. 1998. Limitation of carbon assimilation of intertidal Zostera noltii and Z-marina by desiccation at low tide. Aquat. Bot. 62: 171–176. doi: 10.1016S0304-3770(98)00091-6

Manassa, R. P., T. M. Smith, J. Beardall, M. J. Keough, and P. L. M. Cook. 2017. Capacity of a temperate intertidal seagrass species to tolerate changing environmental conditions: Significance of light and tidal exposure. Ecol. Indic. 81: 578–586. doi: 10.1016/j.ecolind.2017.04.056

Moreno-Marín, F., F. G. Brun, and M. F. Pedersen. 2018. Additive response to multiple environmental stressors in the seagrass Zostera marina L. Limnol. Oceanogr. 63: 1528–1544. doi: 10.1002/lno.10789

Olesen, B., and K. Sand-Jensen. 1993. Seasonal acclimatization of eelgrass Zostera marina growth to light. Mar. Ecol. Prog. Ser. 94: 91–91. doi: 10.3354/meps094091

Orth, R. J. and others. 2020. Restoration of seagrass habitat leads to rapid recovery of coastal ecosystem services. Sci. Adv. 6: eabc6434. doi: 10.1126/sciadv.abc6434

Papathanasiou, V., G. Kariofillidou, P. Malea, and S. Orfanidis. 2020. Effects of air exposure on desiccation and photosynthetic performance of Cymodocea nodosa with and without epiphytes and Ulva rigida in comparison, under laboratory conditions. Mar. Environ. Res. 158: 104948. doi: 10.1016/j.marenvres.2020.104948

Park, S. R., S. Kim, Y. K. Kim, C. K. Kang, and K. S. Lee. 2016. Photoacclimatory Responses of Zostera marina in the Intertidal and Subtidal Zones. PLoS One. 11: e0156214. doi: 10.1371/journal.pone.0156214

Park, S. R., K. Moon, S. H. Kim, and K.-S. Lee. 2021. Growth and Photoacclimation Strategies of Three Zostera Species Along a Vertical Gradient: Implications for Seagrass Zonation Patterns. Front. Mar. Sci. 8: 594779. doi: 10.3389/fmars.2021.594779

Pérez-Lloréns, J., S. Strother, and F. Niell. 1994. Species differences in short-term pigment levels in four Australian seagrasses in response to desiccation and rehydration. Bot. Mar. 37: 91–96. doi: 10.1515botm.1994.37.1.91

Petrou, K., I. Jimenez-Denness, K. Chartrand, C. McCormack, M. Rasheed, and P. J. Ralph. 2013. Seasonal heterogeneity in the photophysiological response to air exposure in two tropical intertidal seagrass species. Mar. Ecol. Prog. Ser. 482: 93–106. doi: 10.3354/meps10229

Piercy, C. D., B. R. Charbonneau, E. R. Russ, and T. M. Swannack. 2023. Examining the commonalities and knowledge gaps in coastal zone vegetation simulation models. Earth Surf. Process. Landf.: 1–25. doi: 10.1002/esp.5565

Poorter, H., N. P. Anten, and L. F. Marcelis. 2013. Physiological mechanisms in plant growth models: do we need a supra-cellular systems biology approach? Plant Cell Environ. 36: 1673–1690. doi: 10.1111/pce.12123

Scalpone, C. R., J. C. Jarvis, J. M. Vasslides, J. M. Testa, and N. K. Ganju. 2020. Simulated Estuary-Wide Response of Seagrass (Zostera marina) to Future Scenarios of Temperature and Sea Level. Front. Mar. Sci. 7: 873. doi:

Seddon, S., and A. C. Cheshire. 2001. Photosynthetic response of Amphibolis antarctica and Posidonia australis to temperature and desiccation using chlorophyll fluorescence. Mar. Ecol. Prog. Ser. 220: 119–130. doi: 10.3354/meps220119

Shafer, D. J., T. D. Sherman, and S. Wyllie-Echeverria. 2007. Do desiccation tolerances control the vertical distribution of intertidal seagrasses? Aquat. Bot. 87: 161–166. doi: 10.1016/j.aquabot.2007.04.003

Silva, J., R. Santos, M. L. Calleja, and C. M. Duarte. 2005. Submerged versus air-exposed intertidal macrophyte productivity: from physiological to community-level assessments. J. Exp. Mar. Biol. Ecol. 317: 87–95. doi: 10.1016/j.jembe.2004.11.010

Simpson, M. J., A. P. Browning, D. J. Warne, O. J. Maclaren, and R. E. Baker. 2022. Parameter identifiability and model selection for sigmoid population growth models. J. Theor. Biol. 535: 110998. doi: 10.1016/j.jtbi.2021.110998

Suykerbuyk, W. and others. 2018. Living in the intertidal: desiccation and shading reduce seagrass growth, but high salinity or population of origin have no additional effect. PeerJ. 6: e5234. doi: 10.7717/peerj.5234

Tanaka, Y., and M. Nakaoka. 2004. Emergence stress and morphological constraints affect the species distribution and growth of subtropical intertidal seagrasses. Mar. Ecol. Prog. Ser. 284: 117–131. doi: 10.3354/meps284117

Tian, R. C. 2006. Toward standard parameterizations in marine biological modeling. Ecol. Model. 193: 363–386. doi: 10.1016/j.ecolmodel.2005.09.003

Toublanc, F., I. Brenon, T. Coulombier, and O. Le Moine. 2015. Fortnightly tidal asymmetry inversions and perspectives on sediment dynamics in a macrotidal estuary (Charente, France). Cont. Shelf Res. 94: 42–54. doi: 10.1016/j.csr.2014.12.009

Turschwell, M. P. and others. 2022. Interactive effects of multiple stressors vary with consumer interactions, stressor dynamics and magnitude. Ecol. Lett. 25: 1483–1496. doi: 10.1111/ele.14013

Vieira, V. M. N. C. S., I. E. Lopes, and J. C. Creed. 2018. The biomass–density relationship in seagrasses and its use as an ecological indicator. BMC Ecology. 18: 44. doi: 10.1186/s12898-018-0200-1

Wang, X., J. Bai, J. Yan, B. Cui, and D. Shao. 2022. How Turbidity Mediates the Combined Effects of Nutrient Enrichment and Herbivory on Seagrass Ecosystems. Front. Mar. Sci. 9: 787041. doi: 10.3389/fmars.2022.787041

Wang, X., J. Yan, J. Bai, D. Shao, and B. Cui. 2021. Effects of interactions between macroalgae and seagrass on the distribution of macrobenthic invertebrate communities at the Yellow River Estuary, China. Mar. Pollut. Bull. 164: 112057. doi: 10.1016/j.marpolbul.2021.112057

Waycott, M. and others. 2009. Accelerating loss of seagrasses across the globe threatens coastal ecosystems. Proc. Natl. Acad. Sci. U.S.A. 106: 12377–12381. doi: 10.1073pnas.0905620106

Wuthirak, T., R. Kongnual, and P. Buapet. 2016. Desiccation tolerance and underlying mechanisms for the recovery of the photosynthetic efficiency in the tropical intertidal seagrasses Halophila ovalis and Thalassia hemprichii. Bot. Mar. 59: 387–396. doi: doi:10.1515/bot-2016-0052

Zhang, X. and others. 2019. A unique meadow of the marine angiosperm Zostera japonica, covering a large area in the turbid intertidal Yellow River Delta, China. Sci. Total. Environ. 686: 118–130. doi: 10.1016/j.scitotenv.2019.05.320

Zhou, Q., Y. Ke, X. Wang, J. Bai, D. Zhou, and X. Li. 2022. Developing seagrass index for long term monitoring of Zostera japonica seagrass bed: A case study in Yellow River Delta, China. ISPRS J. Photogramm. 194: 286–301. doi: 10.1016j.isprsjprs.2022.10.011

