## Supplementary Material for "Trade-off between light deprivation and desiccation in intertidal seagrasses due to periodic tidal inundation and exposure: insights from a data-calibrated model"

**Table S1**

Desiccation coefficient (*k*) and its relationship with air temperature for two seagrass species were derived based on data from Seddon and Cheshire (2001). Desiccation coefficients for individual air temperatures were found from the fit of Eq. (7) to the data shown in Fig. S1 for *Posidonia australis* and Fig. S2 for *Amphibolis antarctica*, and are written here as mean ± SD. The linear model *k*(*T*) was obtained from fitting Eq. (8) to mean values of the desiccation coefficients estimated at individual air temperatures, as shown in Fig. S3.

| Seagrass species | Air temperature, *T* (℃) | Desiccation coefficient, *k* (d^-1^) | Equation |
| --- | --- | --- | --- |
| *Posidonia australis* | 18 | 10.8±0.5 | *k*(*T*)=0.5(*T*-20)+11.6  (R^2^=0.98) |
|  | 24 | 12.9±0.5 |  |
|  | 28 | 15.6±0.9 |  |
|  | 32 | 16.8±0.7 |  |
| *Amphibolis antarctica* | 18 | 8.2±0.1 | *k*(*T*)=0.4(*T-*20)+9.9  (R^2^=0.91) |
|  | 24 | 10.6±0.2 |  |
|  | 28 | 14.1±0.6 |  |
|  | 32 | 14.0±0.7 |  |

**Table S2**

Parameters for $f_{\mathrm{RWC}}\left( \mathrm{RWC} \right)$ were found from the fit of Eq. (10) to the data from Jiang et al. (2014) shown in Fig. S6 for *Thalassia hemprichii* and *T. acoroides*, and the fit of Eq. (11) to the data from Shafer et al. (2007) shown in Fig. S7 for *Zostera japonica* and *Z. marina*.

| Model form of $f_{\mathrm{RWC}}\left( \mathrm{RWC} \right)$ | Seagrass species | Model parameters | | | | |
| --- | --- | --- | --- | --- | --- | --- |
|  |  | Air temperature, *T* (℃) | $\mathrm{RWC}_{\mathrm{crit}}$ | $\mathrm{RWC}_{k}$ | *R* | $\mathrm{RWC}_{h}$ |
| Hyperbolic tangent model | *Thalassia hemprichii* | 24 | 0.01 | 0.2 | - | - |
|  |  | 32 | 0.05 | 0.1 | - | - |
|  | *Thalassia acoroide* | 24 | 0 | 0.3 | - | - |
|  |  | 32 | 0 | 0.3 | - | - |
| Sigmoidal curve model | *Zostera japonica* | 22 | - | - | 15.0 | 0.4 |
|  | *Zostera marina* | 22 | - | - | 10.8 | 0.4 |

**Table S3**

Baseline and testing scenarios for the model application as described in Section 2.4.2. *^a^* Tidal range scenario refers to the seagrasses subjected to varying ranges of high and low tides, parameterized by the amplitudes of the M2 and S2 tidal constituents. Parameterisation of the microtidal scenario is detailed in Table 2, representing a tidal range of 1.6 m. For the mesotidal scenario, the tidal amplitude is 1.0 m and 0.3 m for the M2 and S2 tidal constituents, respectively, yielding an overall tidal range of 2.6 m (Azevedo et al. 2017), while for the macrotidal scenario, these values are 1.8 m and 0.6 m, respectively, yielding an overall tidal range of 4.8 m (Toublanc et al. 2015). *^b^* For scenarios in this table with a meadow elevation differing from 0 m, the lowest and highest meadow elevations tested are given in brackets under the categorical label “Subtidal”, “Intertidal” or “Supratidal” (see Section 2.4.2 for further details).

| Scenarios | Tidal range*^a^* | Meadow elevation*^b^*  ($Z_{b}$, m) | Turbidity  ($K_{d}$, m^-1^) | Growth function | Interactive factor  formulation | $f_{\mathrm{RWC}}\left( \mathrm{RWC} \right)$ |
| --- | --- | --- | --- | --- | --- | --- |
| Baseline | Microtidal | 0 | 0.05 | $\mu(, \mathrm{RWC})$ | Multiplicative | Sigmoidal curve model |
| Group $I$ | Microtidal | 0 | 0.05 | $\mu(I)$ | Not applicable | Not applicable |
|  | Microtidal | 0 | 0.05 | $\mu(I, \mathrm{RWC})$ | Multiplicative | Linear model |
|  | Microtidal | 0 | 0.05 | $\mu(I, \mathrm{RWC})$ | Multiplicative | Hyperbolic tangent model |
|  | Microtidal | 0 | 0.05 | $\mu(I, \mathrm{RWC})$ | Law of the minimum | Sigmoidal model |
|  | Microtidal | 0 | 0.05 | $\mu(I, \mathrm{RWC})$ | Law of the minimum | Linear model |
|  | Microtidal | 0 | 0.05 | $\mu(I, \mathrm{RWC})$ | Law of the minimum | Hyperbolic tangent model |
| Group $\mathrm{II}$ | Microtidal | Subtidal  (-3.0~-0.8) | 0.05 | $\mu(I, \mathrm{RWC})$ | Multiplicative | Sigmoidal curve model |
|  | Microtidal | Intertidal  (-0.8~0.8) | 0.05 | $\mu(I, \mathrm{RWC})$ | Multiplicative | Sigmoidal curve model |
|  | Microtidal | Supratidal  (0.8~3.0) | 0.05 | $\mu(I, \mathrm{RWC})$ | Multiplicative | Sigmoidal curve model |
|  | Microtidal | Subtidal  (-3.0~-0.8) | 0.5 | $\mu(I, \mathrm{RWC})$ | Multiplicative | Sigmoidal curve model |
|  | Microtidal | Intertidal  (-0.8~0.8) | 0.5 | $\mu(I, \mathrm{RWC})$ | Multiplicative | Sigmoidal curve model |
|  | Microtidal | Supratidal  (0.8~3.0) | 0.5 | $\mu(I, \mathrm{RWC})$ | Multiplicative | Sigmoidal curve model |
|  | Microtidal | Subtidal  (-3.0~-0.8) | 1 | $\mu(I, \mathrm{RWC})$ | Multiplicative | Sigmoidal curve model |
|  | Microtidal | Intertidal  (-0.8~0.8) | 1 | $\mu(I, \mathrm{RWC})$ | Multiplicative | Sigmoidal curve model |
|  | Microtidal | Supratidal  (0.8~3.0) | 1 | $\mu(I, \mathrm{RWC})$ | Multiplicative | Sigmoidal curve model |
|  | Microtidal | Subtidal  (-3.0~-0.8) | 1.5 | $\mu(I, \mathrm{RWC})$ | Multiplicative | Sigmoidal curve model |
|  | Microtidal | Intertidal  (-0.8~0.8) | 1.5 | $\mu(I, \mathrm{RWC})$ | Multiplicative | Sigmoidal curve model |
|  | Microtidal | Supratidal  (0.8~3.0) | 1.5 | $\mu(I, \mathrm{RWC})$ | Multiplicative | Sigmoidal curve model |
|  | Microtidal | Subtidal  (-3.0~-0.8) | 2 | $\mu(I, \mathrm{RWC})$ | Multiplicative | Sigmoidal curve model |
|  | Microtidal | Intertidal  (-0.8~0.8) | 2 | $\mu(I, \mathrm{RWC})$ | Multiplicative | Sigmoidal curve model |
|  | Microtidal | Supratidal  (0.8~3.0) | 2 | $\mu(I, \mathrm{RWC})$ | Multiplicative | Sigmoidal curve model |
| Group $\mathrm{III}$ | Mesotidal | Subtidal  (-3.0~-1.3) | 0.05 | $\mu(I, \mathrm{RWC})$ | Multiplicative | Sigmoidal curve model |
|  | Mesotidal | Intertidal  (-1.3~1.3) | 0.05 | $\mu(I, \mathrm{RWC})$ | Multiplicative | Sigmoidal curve model |
|  | Mesotidal | Supratidal  (1.3~3.0) | 0.05 | $\mu(I, \mathrm{RWC})$ | Multiplicative | Sigmoidal curve model |
|  | Mesotidal | Subtidal  (-3.0~-1.3) | 0.5 | $\mu(I, \mathrm{RWC})$ | Multiplicative | Sigmoidal curve model |
|  | Mesotidal | Intertidal  (-1.3~1.3) | 0.5 | $\mu(I, \mathrm{RWC})$ | Multiplicative | Sigmoidal curve model |
|  | Mesotidal | Supratidal  (1.3~3.0) | 0.5 | $\mu(I, \mathrm{RWC})$ | Multiplicative | Sigmoidal curve model |
|  | Mesotidal | Subtidal  (-3.0~-1.3) | 1 | $\mu(I, \mathrm{RWC})$ | Multiplicative | Sigmoidal curve model |
|  | Mesotidal | Intertidal  (-1.3~1.3) | 1 | $\mu(I, \mathrm{RWC})$ | Multiplicative | Sigmoidal curve model |
|  | Mesotidal | Supratidal  (1.3~3.0) | 1 | $\mu(I, \mathrm{RWC})$ | Multiplicative | Sigmoidal curve model |
|  | Mesotidal | Subtidal  (-3.0~-1.3) | 1.5 | $\mu(I, \mathrm{RWC})$ | Multiplicative | Sigmoidal curve model |
|  | Mesotidal | Intertidal  (-1.3~1.3) | 1.5 | $\mu(I, \mathrm{RWC})$ | Multiplicative | Sigmoidal curve model |
|  | Mesotidal | Supratidal  (1.3~3.0) | 1.5 | $\mu(I, \mathrm{RWC})$ | Multiplicative | Sigmoidal curve model |
|  | Mesotidal | Subtidal  (-3.0~-1.3) | 2 | $\mu(I, \mathrm{RWC})$ | Multiplicative | Sigmoidal curve model |
|  | Mesotidal | Intertidal  (-1.3~1.3) | 2 | $\mu(I, \mathrm{RWC})$ | Multiplicative | Sigmoidal curve model |
|  | Mesotidal | Supratidal  (1.3~3.0) | 2 | $\mu(I, \mathrm{RWC})$ | Multiplicative | Sigmoidal curve model |
| Group $\mathrm{IV}$ | Macrotidal | Subtidal  (-3.0~-2.4) | 0.05 | $\mu(I, \mathrm{RWC})$ | Multiplicative | Sigmoidal curve model |
|  | Macrotidal | Intertidal  (-2.4~2.4) | 0.05 | $\mu(I, \mathrm{RWC})$ | Multiplicative | Sigmoidal curve model |
|  | Macrotidal | Supratidal  (2.4~3.0) | 0.05 | $\mu(I, \mathrm{RWC})$ | Multiplicative | Sigmoidal curve model |
|  | Macrotidal | Subtidal  (-3.0~-2.4) | 0.5 | $\mu(I, \mathrm{RWC})$ | Multiplicative | Sigmoidal curve model |
|  | Macrotidal | Intertidal  (-2.4~2.4) | 0.5 | $\mu(I, \mathrm{RWC})$ | Multiplicative | Sigmoidal curve model |
|  | Macrotidal | Supratidal  (2.4~3.0) | 0.5 | $\mu(I, \mathrm{RWC})$ | Multiplicative | Sigmoidal curve model |
|  | Macrotidal | Subtidal  (-3.0~-2.4) | 1 | $\mu(I, \mathrm{RWC})$ | Multiplicative | Sigmoidal curve model |
|  | Macrotidal | Intertidal  (-2.4~2.4) | 1 | $\mu(I, \mathrm{RWC})$ | Multiplicative | Sigmoidal curve model |
|  | Macrotidal | Supratidal  (2.4~3.0) | 1 | $\mu(I, \mathrm{RWC})$ | Multiplicative | Sigmoidal curve model |
|  | Macrotidal | Subtidal  (-3.0~-2.4) | 1.5 | $\mu(I, \mathrm{RWC})$ | Multiplicative | Sigmoidal curve model |
|  | Macrotidal | Intertidal  (-2.4~2.4) | 1.5 | $\mu(I, \mathrm{RWC})$ | Multiplicative | Sigmoidal curve model |
|  | Macrotidal | Supratidal  (2.4~3.0) | 1.5 | $\mu(I, \mathrm{RWC})$ | Multiplicative | Sigmoidal curve model |
|  | Macrotidal | Subtidal  (-3.0~-2.4) | 2 | $\mu(I, \mathrm{RWC})$ | Multiplicative | Sigmoidal curve model |
|  | Macrotidal | Intertidal  (-2.4~2.4) | 2 | $\mu(I, \mathrm{RWC})$ | Multiplicative | Sigmoidal curve model |
|  | Macrotidal | Supratidal  (2.4~3.0) | 2 | $\mu(I, \mathrm{RWC})$ | Multiplicative | Sigmoidal curve model |

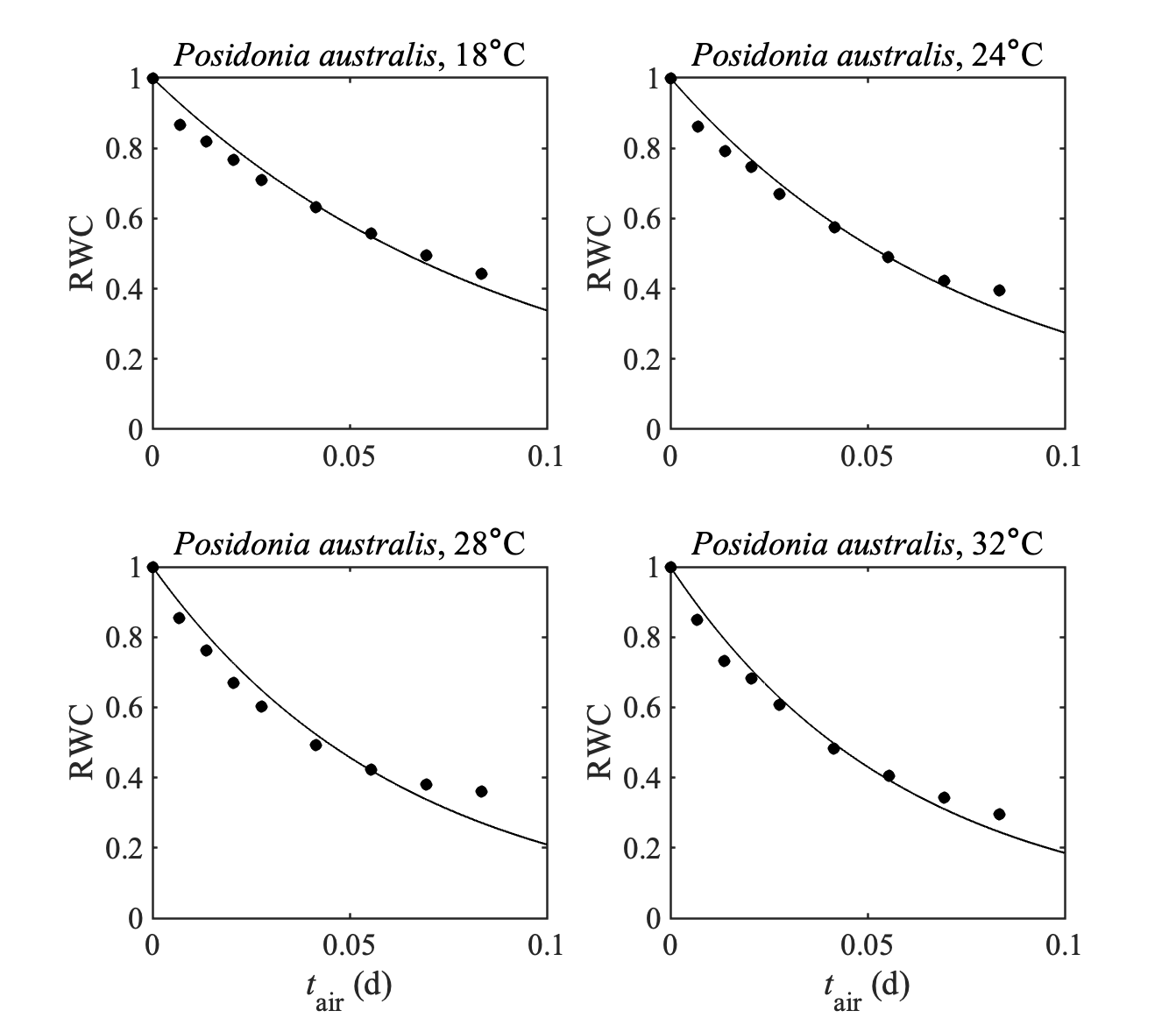

**Fig. S1.** Data (dots) and fitted models (curved lines, Eq. (7)) for the relationship between RWC and air-exposure duration $t_{\mathrm{air}}$ at four air temperatures for the seagrass species *P. australis*. Data obtained from Seddon and Cheshire (2001). The model-data fit was reasonable for all four air temperatures (R^2^ > 0.95 and *p* < 0.001). Parameter estimates (mean ± SD) for *k*(*T*) obtained from these model-data fits are given in Table S1.

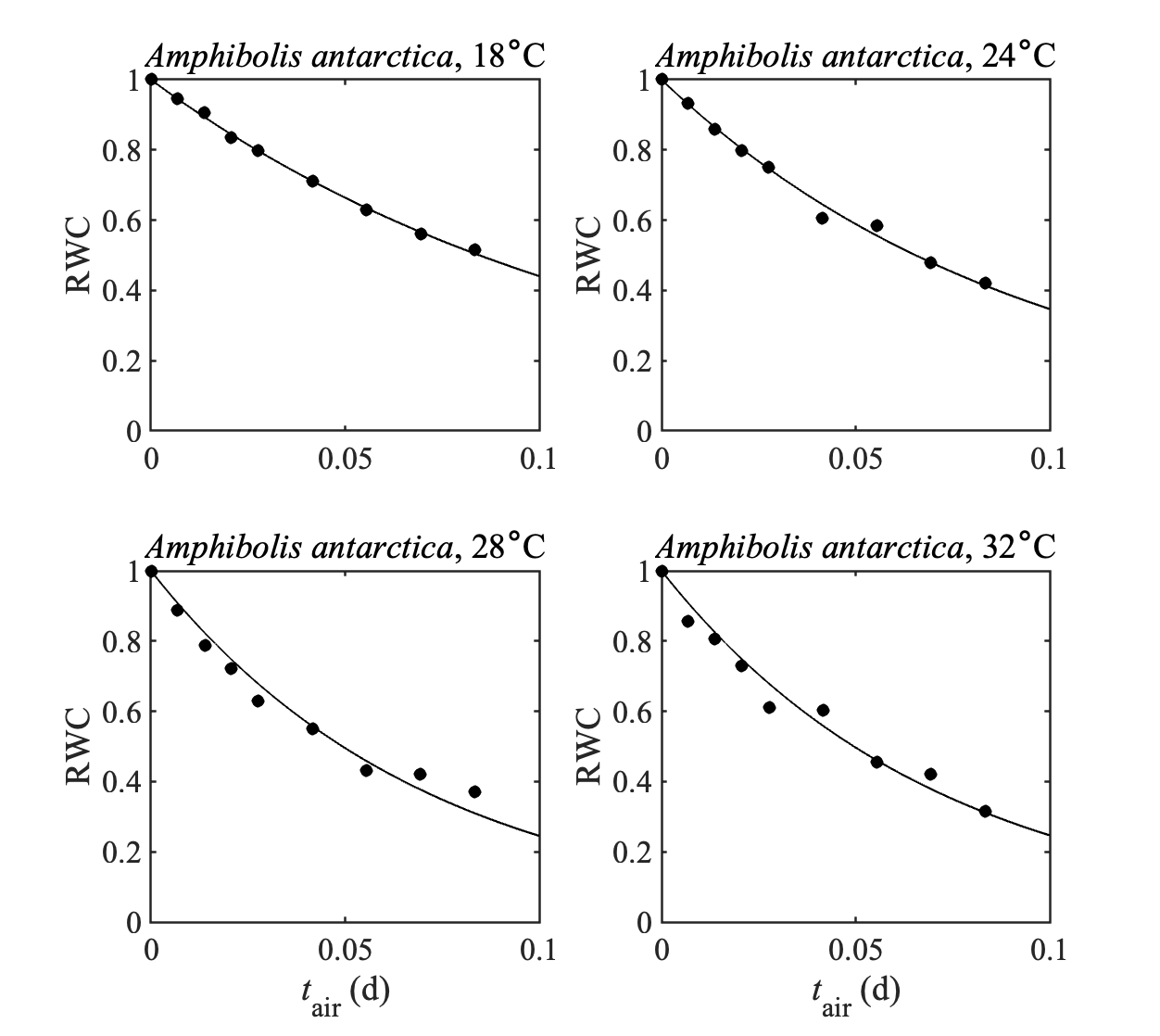

**Fig. S2.** Data (dots) and fitted models (curved lines, Eq. (7)) for the relationship between RWC and air- exposure duration $t_{\mathrm{air}}$ at four air temperatures for the seagrass species *A. antarctica*. Data obtained from Seddon and Cheshire (2001). The model-data fit was reasonable for all four air temperatures (R^2^ > 0.95 and *p* < 0.001). Parameter estimates (mean ± SD) for *k*(*T*) obtained from these model-data fits are given in Table S1.

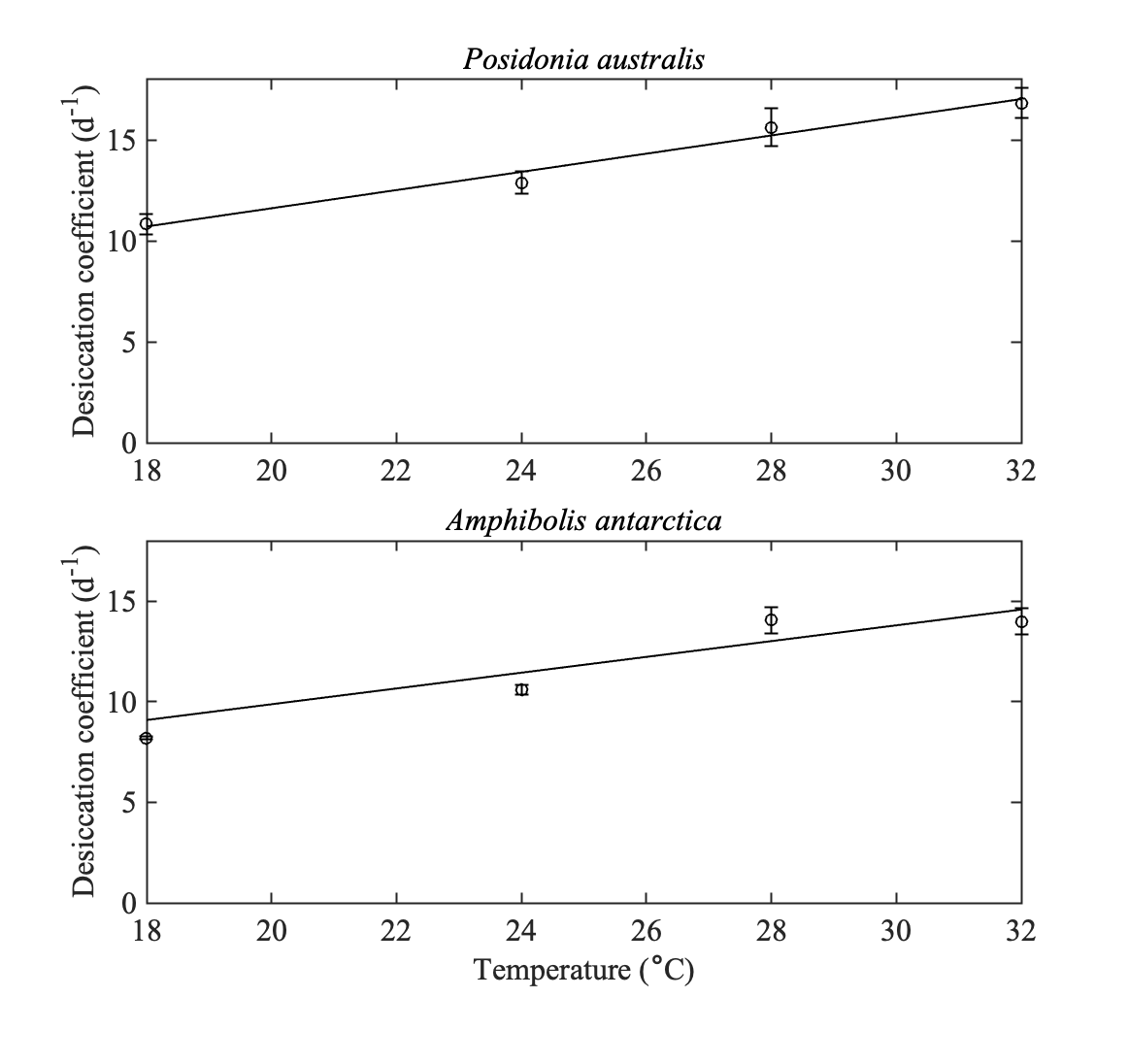

**Fig. S3.** Estimated desiccation coefficients at individual air temperatures (dots and error bars) and fitted models (lines, Eq. (8)) for the relationship *k*(*T*) between desiccation coefficient and air temperature for the seagrass species *P. australis* and *A. antarctica*. Estimated desiccation coefficients at individual air temperatures were obtained from model-data fits in Fig. S1 and S2 to data from Seddon and Cheshire (2001). The model-data fit was reasonable for both species (R^2^ > 0.9 and *p* < 0.05). The fitted linear relationships *k*(*T*) are given in Table S1.

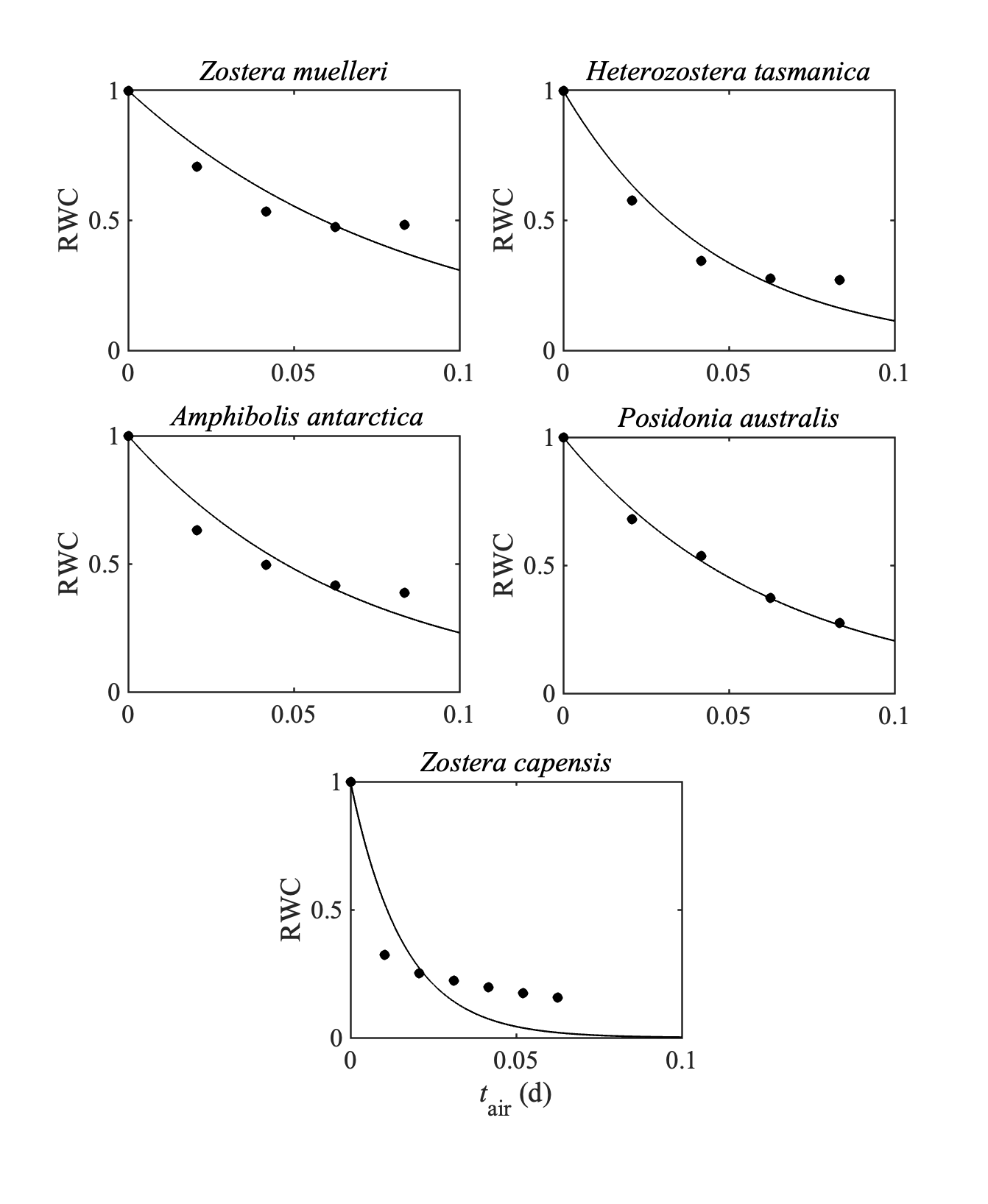

**Fig. S4.** Data (dots) and fitted models (curved lines, Eq. (7)) for the relationship between RWC and air- exposure duration $t_{\mathrm{air}}$ for the seagrass species *Z. muelleri*, *H. tasmanica, A. antarctica*, *P. australis* (data were obtained from Pérez-Lloréns et al. (1994)) and *Z. capensis* (data was obtained from Adams and Bate (1994)). The model-data fit was reasonable for all species (R^2^ > 0.85 and *p* < 0.01) except for *Z. capensis* (R^2^ > 0.8 and *p* < 0.01). Parameter estimates (mean) obtained from these model-data fits are given in Table 1.

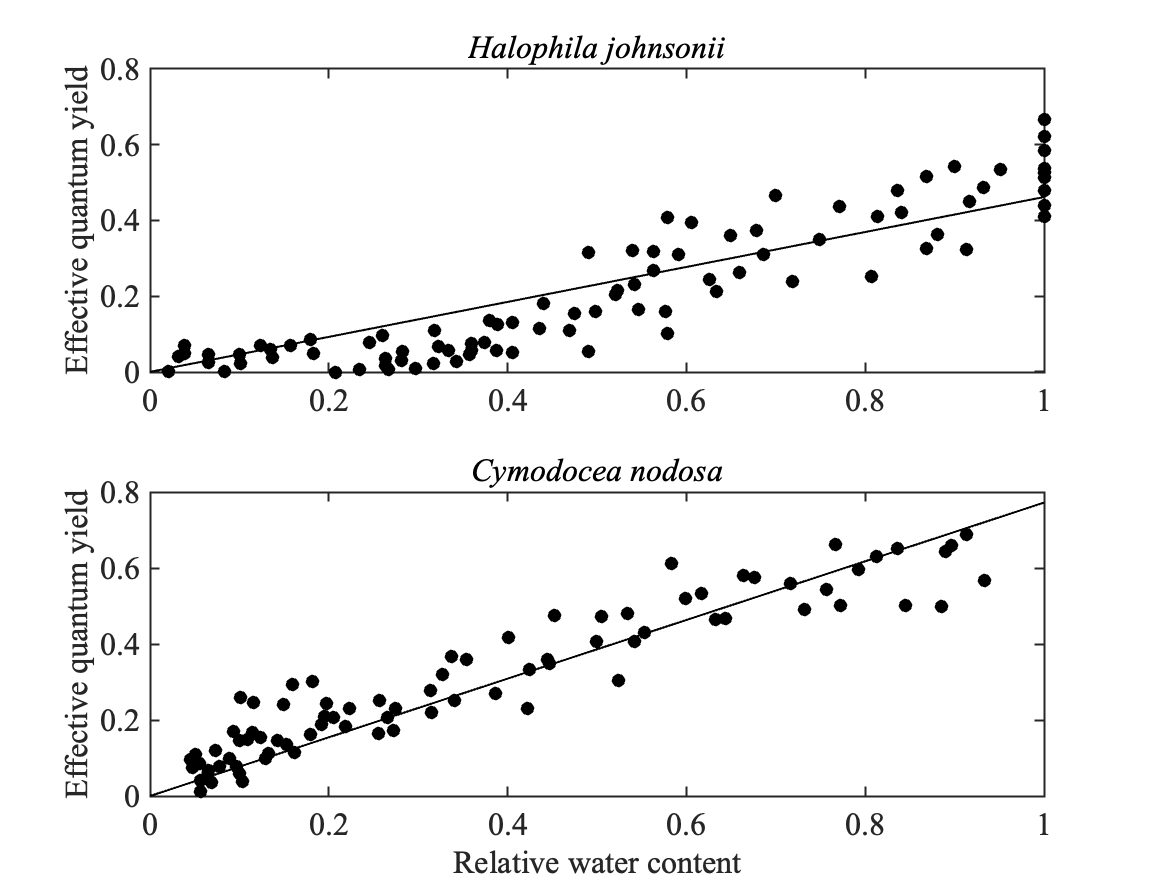

**Fig. S5.** Relationship between effective quantum yield and relative water content for the seagrass species *H. johnsonii* and *C. nodosa.* A linear function (Eq. (9)) provided a good fit to the data from Kahn and Durako (2009) and Papathanasiou et al. (2020) for *H. johnsonii* (R^2^=0.79, *p* < 0.001) and *C. nodosa* (R^2^=0.85, *p* < 0.001), respectively.

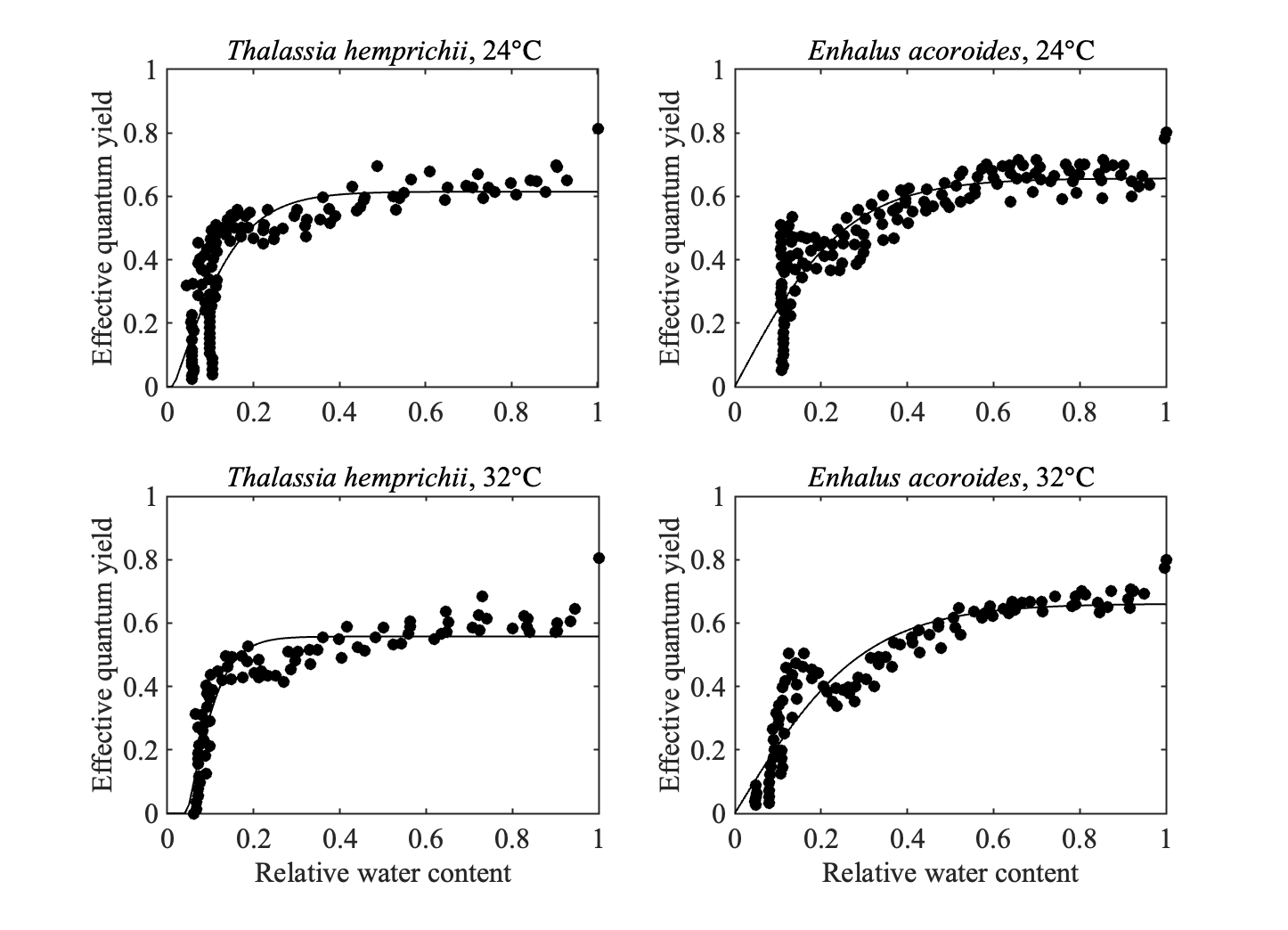

**Fig. S6.** Data (dots) and fitted model (Eq. (10), assuming RWC_crit_ ≥ 0, line) for the relationship between effective quantum yield and RWC for intertidal *T. hemprichii* and *E. acoroides* at different air temperatures. Data were obtained from Jiang et al. (2014)*.* The model-data fit was more reasonable when the air temperature is 32 ℃ (R^2^>0.8 and *p* < 0.001 for both species) than when the air temperature is 24 ℃ (R^2^>0.7 and *p* < 0.001 for both species). Parameter estimates (mean) obtained from these model-data fits are given in Table S2.

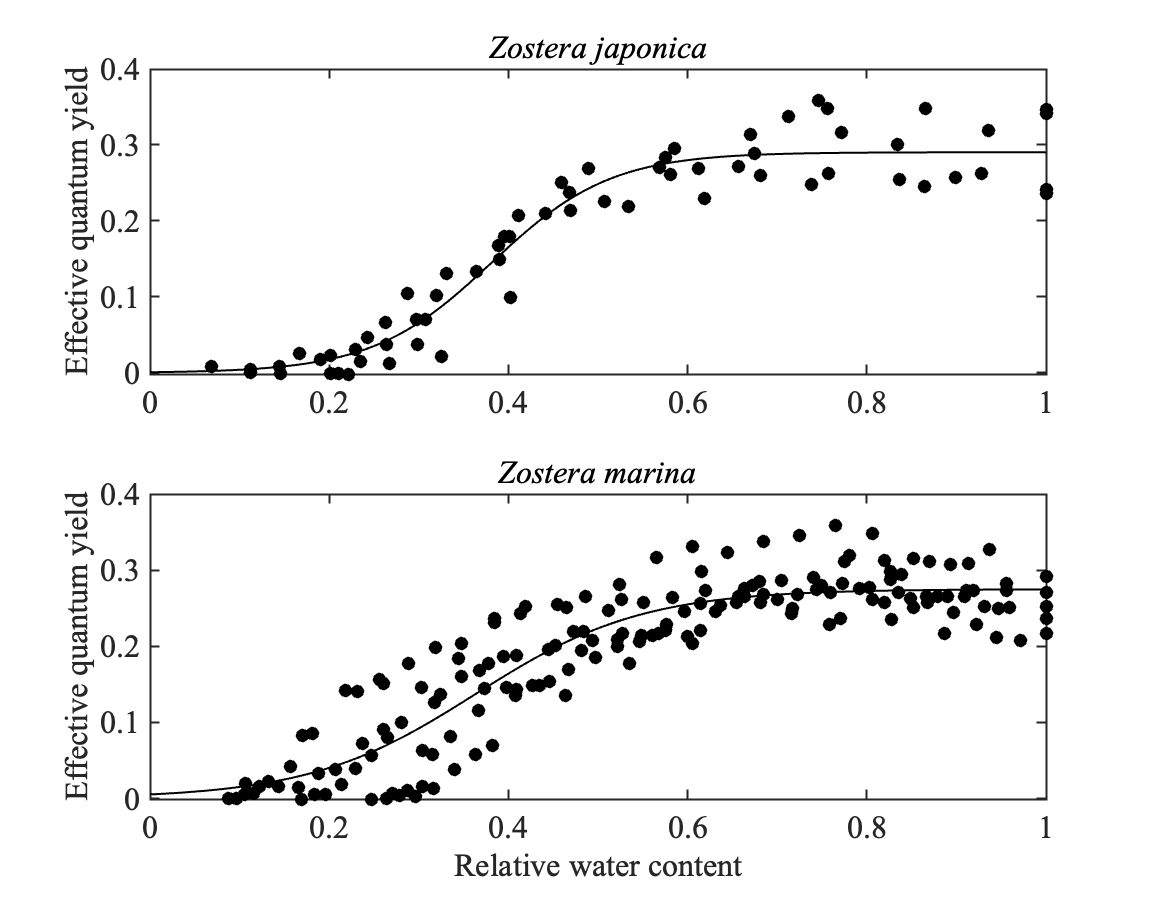

**Fig. S7.** Relationship between effective quantum yield and relative water content for the seagrass species *Z. japonica* and *Z.* *marina.* The fitted function (Eq. (11)) provided a good fit to the data from Shafer et al. (2007) for *Z. japonica* (R^2^=0.93, *p* < 0.001) and *Z.* *marina* (R^2^=0.82, *p* < 0.001), respectively. Parameter estimates (mean) obtained from these model-data fits are given in Table S2.

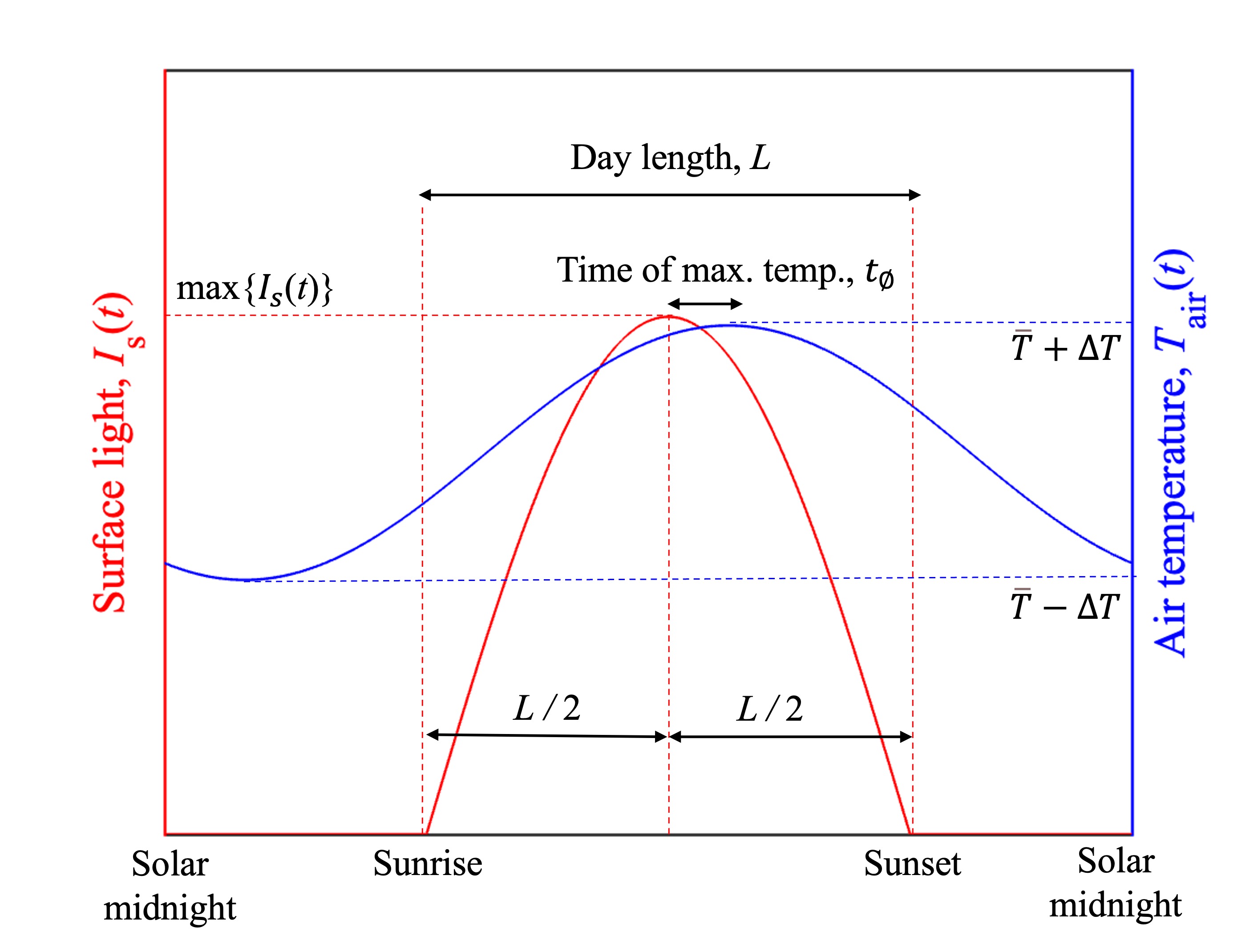

**Fig. S8.** The daily cycle of surface light $I_{s}(t)$ (red line), and air temperature $T_{\mathrm{air}}(t)$ (blue) assumed in the minimum realistic models of surface light and air temperature described in Section 2.3. The figure is adapted from Adams et al. (2020).

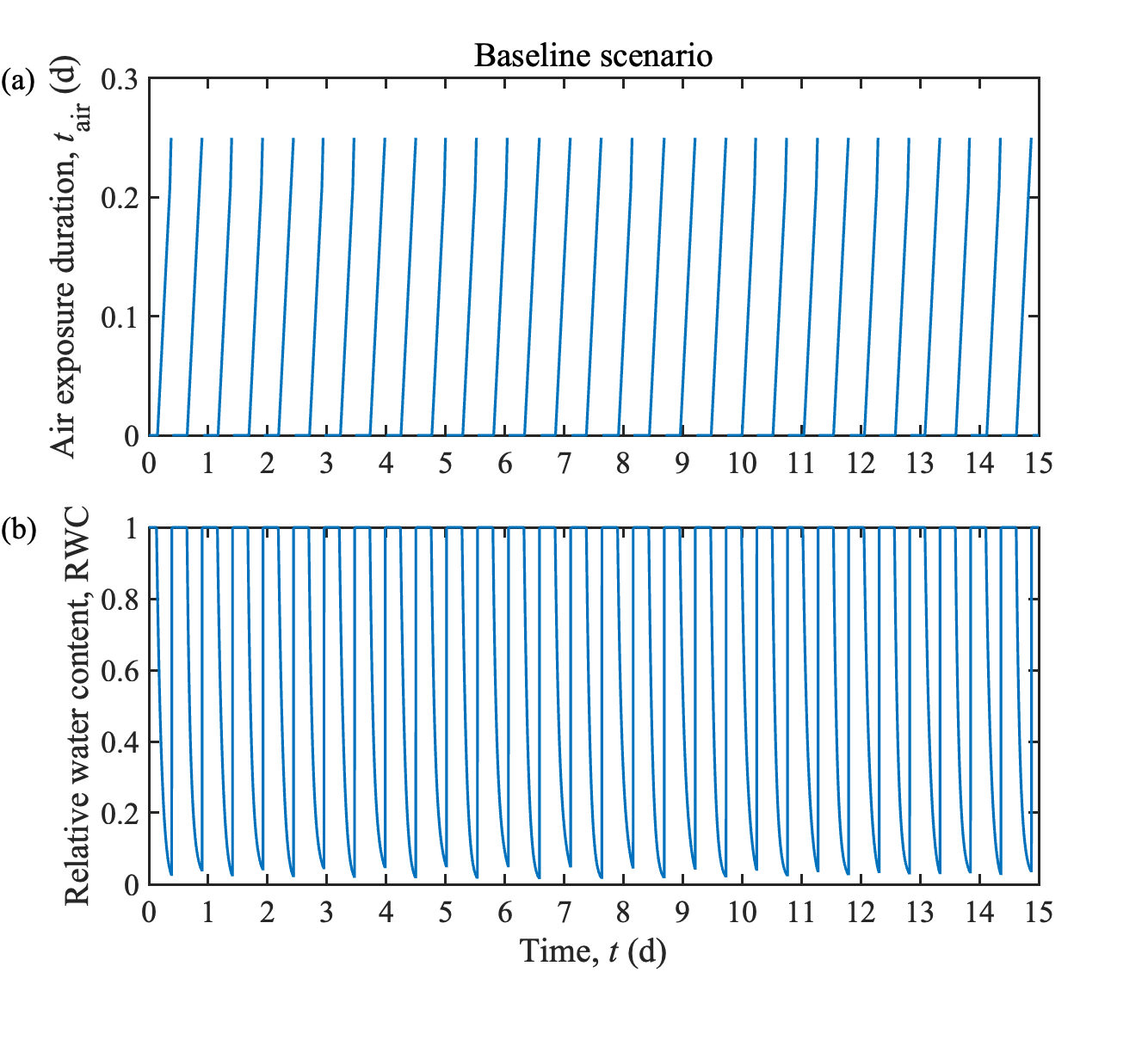

**Fig. S9.** The results of modelled (a) air-exposure duration, $t_{air}$, and (b) RWC within 15 d for the Baseline scenario (Table 2) of simulating intertidal seagrass.

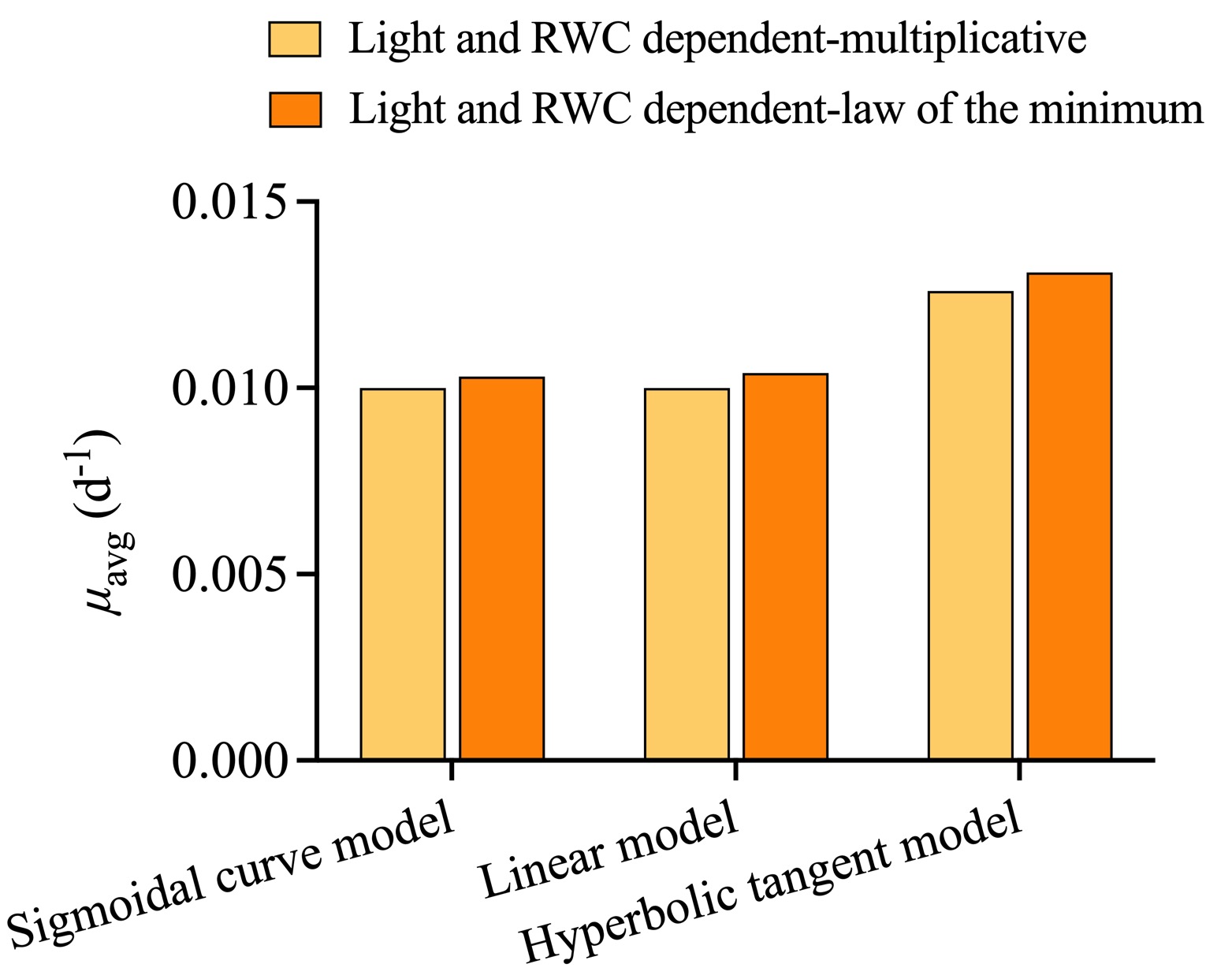

**Fig. S10.** The results of modelled average growth rate $(\mu_{\mathrm{avg}})$ during the simulation period of 15 d for light and RWC dependent growth rate followed the multiplicative or law of the minimum formulation with the three types of $f_{\mathrm{RWC}}\left( \mathrm{RWC} \right)$ defined in Eq. (9)-(11) at lower irradiances (10 mol m^-2^ d^-1^).

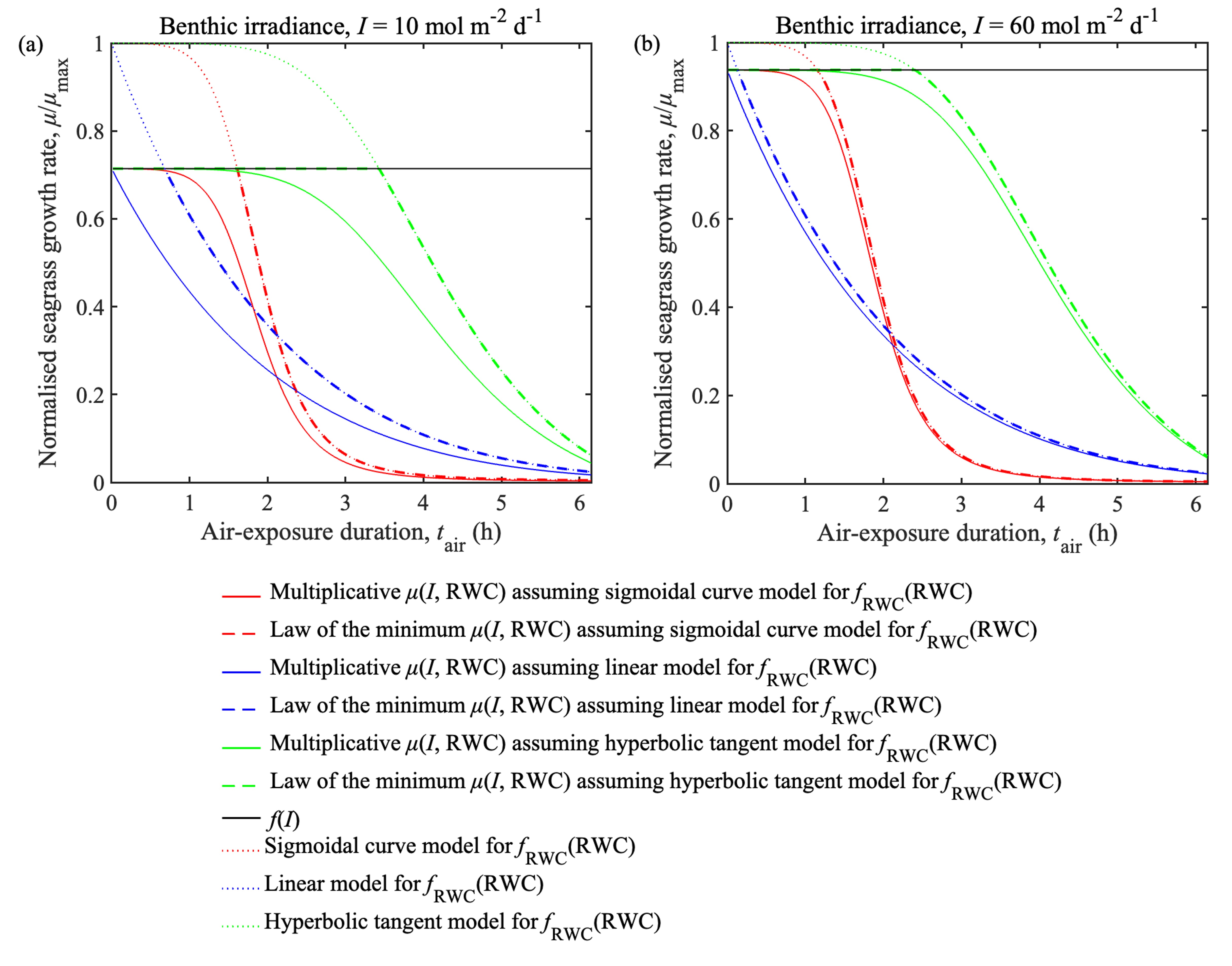

**Fig. S11.** The relationship between the normalized growth rate ($\mu/\mu_{\max}$) and air-exposure duration ($t_{\mathrm{air}}$) for two seagrass growth functions with three types of $f_{\mathrm{RWC}}\left( \mathrm{RWC} \right)$ under two light irradiance conditions: (a) 10 mol m^-2^ d^-1^ and (b) 60 mol m^-2^ d^-1^.

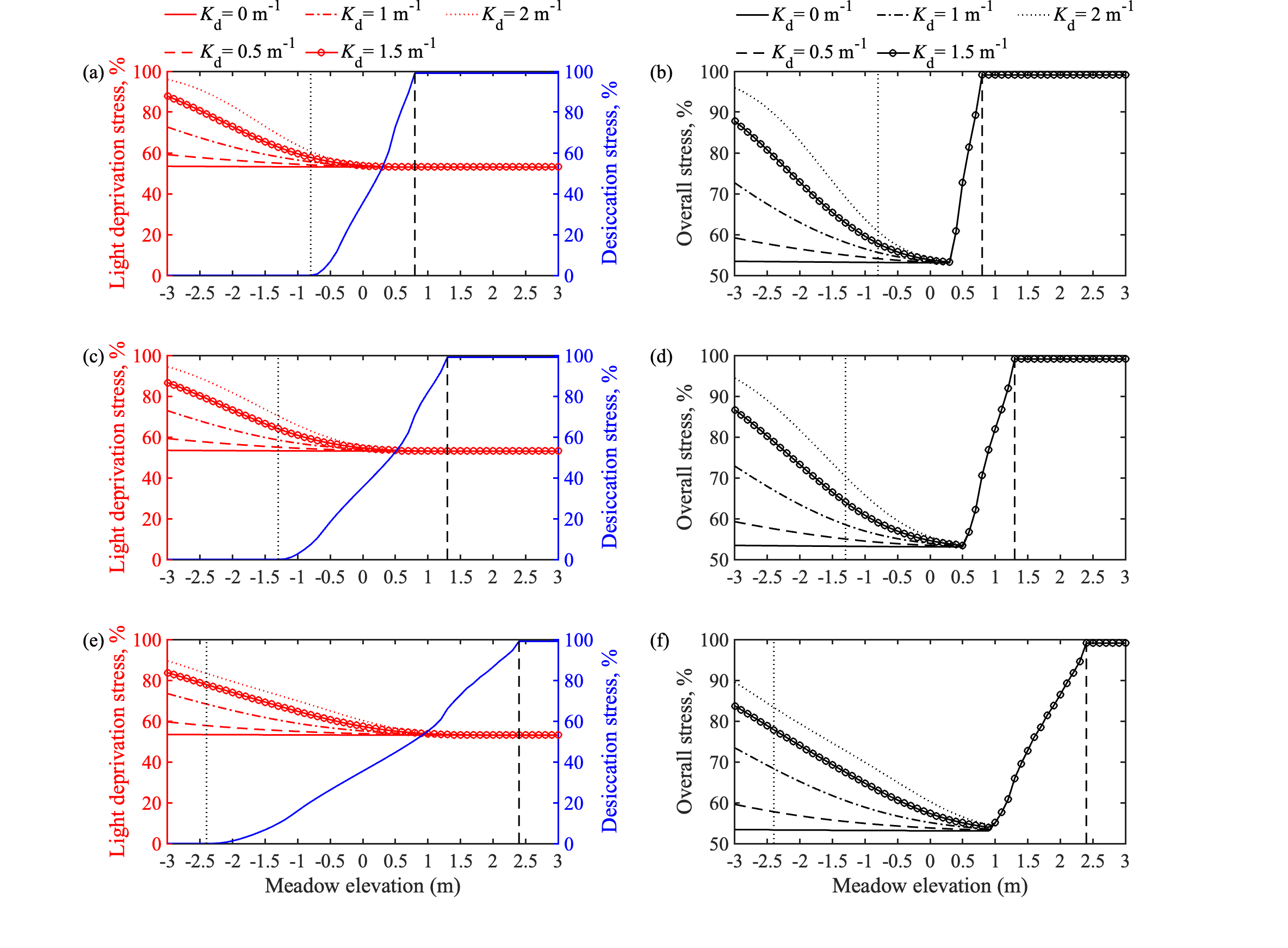

**Fig. S12.** The modelled results of light deprivation stress and desiccation stress on temporally averaged seagrass growth with the law of the minimum formulation over 15 d along the vertical depth gradient under different water turbidity conditions and tidal conditions. The separate effects of light deprivation stress and desiccation stress are shown in panels (a, c, e) and the combined effect of both stressors is shown in panels (b, d, f). Predictions are shown for microtidal conditions (a, b), mesotidal conditions (c, d), and macrotidal conditions (e, f). In each of the six panels, the black dotted line is the boundary between subtidal and lower intertidal zones, while the black dashed line is the boundary between upper intertidal and supratidal zones. Meadow elevation $Z_{b}$(m) is relative to mean sea level, and $K_{d}$ (m^-1^) is the light attenuation coefficient of the water column. These plots were constructed using the Baseline and Groups II, III and IV modelling scenarios (Table S3) that assumed the law of the minimum formulation for $\mu\left( I, \mathrm{RWC} \right)$.

**Supplementary Material References**

Adams, J., and G. Bate. 1994. The tolerance to desiccation of the submerged macrophytes Ruppia cirrhosa (Petagna) Grande and Zostera capensis Setchell. *J. Exp. Mar. Biol. Ecol.* **183:** 53-62. doi: 10.1016/0022-0981(94)90156-2

Adams, M. P. and others. 2020. Informing management decisions for ecological networks, using dynamic models calibrated to noisy time-series data. *Eco. Lett.* **23:** 607-619. doi: 10.1111/ele.13465

Azevedo, A., A. I. Lillebø, J. Lencart e Silva, and J. M. Dias. 2017. Intertidal seagrass models: Insights towards the development and implementation of a desiccation module. *Ecol. Model.* **354:** 20-25. doi: 10.1016/j.ecolmodel.2017.03.004

Jiang, Z., X. Huang, J. Zhang, C. Zhou, Z. Lian, and Z. Ni. 2014. The effects of air exposure on the desiccation rate and photosynthetic activity of Thalassia hemprichii and Enhalus acoroides. *Mar. Biol.* **161:** 1051-1061. doi: 10.1007/s00227-014-2398-6

Kahn, A. E., and M. J. Durako. 2009. Photosynthetic tolerances to desiccation of the co-occurring seagrasses Halophila johnsonii and Halophila decipiens. *Aquat. Bot.* **90:** 195-198. doi: 10.1016/j.aquabot.2008.07.003

Papathanasiou, V., G. Kariofillidou, P. Malea, and S. Orfanidis. 2020. Effects of air exposure on desiccation and photosynthetic performance of Cymodocea nodosa with and without epiphytes and Ulva rigida in comparison, under laboratory conditions. *Mar. Environ. Res.* **158:** 104948. doi: 10.1016/j.marenvres.2020.104948

Pérez-Lloréns, J., S. Strother, and F. Niell. 1994. Species differences in short-term pigment levels in four Australian seagrasses in response to desiccation and rehydration. *Bot. Mar.* **37:** 91-96. doi: 10.1515botm.1994.37.1.91

Seddon, S., and A. C. Cheshire. 2001. Photosynthetic response of Amphibolis antarctica and Posidonia australis to temperature and desiccation using chlorophyll fluorescence. *Mar. Ecol. Prog. Ser.* **220:** 119-130. doi: 10.3354/meps220119

Shafer, D. J., T. D. Sherman, and S. Wyllie-Echeverria. 2007. Do desiccation tolerances control the vertical distribution of intertidal seagrasses? *Aquat. Bot.* **87:** 161-166. doi: 10.1016/j.aquabot.2007.04.003

Toublanc, F., I. Brenon, T. Coulombier, and O. Le Moine. 2015. Fortnightly tidal asymmetry inversions and perspectives on sediment dynamics in a macrotidal estuary (Charente, France). *Cont. Shelf Res.* **94:** 42-54. doi: 10.1016/j.csr.2014.12.009
